# A path towards SARS-CoV-2 attenuation: metabolic pressure on CTP synthesis rules the virus evolution

**DOI:** 10.1101/2020.06.20.162933

**Authors:** Zhihua Ou, Christos Ouzounis, Daxi Wang, Wanying Sun, Junhua Li, Weijun Chen, Philippe Marlière, Antoine Danchin

## Abstract

Fighting the COVID-19 epidemic summons deep understanding of the way SARS-CoV-2 taps into its host cell metabolic resources. We describe here the singular metabolic background that creates a bottleneck constraining coronaviruses to evolve towards likely attenuation in the long term. Cytidine triphosphate (CTP) is at the crossroad of the biosynthetic processes that allow the virus to multiply. This is because CTP is in demand for three essential steps. It is a building block of the virus genome, it is required for synthesis of the cytosine-based liponucleotide precursors of the viral envelope and, finally, it is a critical building block of the host transfer RNAs synthesis. The CCA 3’-end of all the transfer RNAs required to translate the RNA genome and further transcripts into the proteins used to build active virus copies is not coded in the human genome. It must be synthesized de novo from CTP and ATP. Furthermore, intermediary metabolism is built on compulsory steps of synthesis and salvage of cytosine-based metabolites via uridine triphosphate (UTP) that keep limiting CTP availability. As a consequence, accidental replication errors tend to replace cytosine by uracil in the genome, unless recombination events allow the sequence to return to its ancestral sequences. We document some of the consequences of this situation in the function of viral proteins. We also highlight and provide a *raison d’être* to viperin, an enzyme of innate antiviral immunity, which synthesizes 3’-deoxy-3′,4’-didehydro-CTP (ddhCTP) as an extremely efficient antiviral nucleotide.

## I INTRODUCTION

The COVID-19 pandemic motivated a deluge of literature investigating the SARS-CoV-2 coronavirus and its epidemic propagation patterns—see (1) and https://pages.semanticscholar.org/coronavirus-research. Detailed compositional and structural analyses both of the virus genome and of the proteins it encodes keep accumulating at a fast pace (https://viralzone.expasy.org/8996). Yet, studies dealing with the way the virus taps into its cell host’s metabolism are scarce. Exploring the nucleotide complement of the virus genome setup in parallel with the corresponding constraints on the proteins it encodes, we investigated here how unique metabolic features impact on the virus’ functions, as we aim at understanding and possibly revealing alleviation of its virulence. A major feature explaining its pathogenicity is that the coronavirus genome mimics the structure of cellular mRNAs, starting with a conventional 5’-end methylated cap (2) and ending up with a 3’-polyadenylated tail. While remarkably apt to create a stealthy contraption, this particular organization makes that the general outline of the viral genome is an obstacle to antiviral therapy. The SARS-CoV-2 genome is so similar to that of the host’s mRNAs that standard interference with the virus expression machinery will often also interfere with that of non-infected cells and be toxic to the host. A second hallmark of the virus is that it is an enveloped virus. Being directly derived from the host, both these features are a handicap when looking for therapeutic solutions. Consequently, these attributes imply drawing resources from the cell’s nucleotide and lipid metabolism. Because the virus had to evolve ways to discriminate its fine attributes from those of the host mRNAs and membranes, it must display idiosyncratic metabolic features that might be used as antiviral targets.

A noteworthy feature shared by envelope construction and RNA synthesis is that both processes rely on cytosine triphosphate (CTP) availability. This prompted us to analyse the consequences of the nucleotide requirement for lipid construction of the viral envelope and the synthesis of its genome, as compared to the host cell’s metabolism that ends up as mRNAs and membranes. We previously discussed how the series of events which begins with copying the virus positive-sense RNA into a general template minus-sense RNA that serves to generate new viral genomes and several individual transcripts of that template (3, 4) is tightly linked to the metabolism of cytosine-containing nucleotides (5). We document here a further singular role of CTP in its mandatory requirement for tRNA maturation into a functional entity, as this impacts availability of a functional translation machinery. It had been noticed that the virus exploits a critical set of pyrimidine-related metabolic pathways to access the pool of ribonucleoside triphosphates needed for the RNA-dependent replication and transcription of the replicated RNA minus strand (6). However, the specific role of CTP was overlooked. In addition to construction of copies of the genome a subset of lipid metabolism—based on cytosine-containing nucleotides—is also recruited to allow for the synthesis of its enveloped capsid. This makes the viral sequence highly sensitive to metabolic details of the cell’s CTP pool synthesis and maintenance, likely to be reflected in the virus evolution as it mutates with a slow general decrease in cytosine nucleotides, attributed at this time to causes that widely differ from what we present here—see e.g. (7). By contrast, functional analysis could help us reveal unexpected key functions of the virus, marked by a divergent trend in the local content of cytosine nucleotides. For example, if the presence of a subset of amino acids—e.g. proline residues—in viral proteins was essential for key functions required for long term survival, then a local increase in cytosine residues would expand the evolutionary landscape of the virus. This type of local bias has indeed been highlighted in a previously discovered feature of coronavirus adaptation to the human host: in SARS-CoV-1 a GC-rich critical sequence of the virus spike protein—a leucine to alanine mutation derived from a GC local enrichment, with a UUA codon changed into GCA—displayed positive selection in the course of evolution of the SARS disease in 2003 (8). Here we depict first, with emphasis on SARS-CoV-2, the details of cytosine-based metabolism that must be retained as a unique coordinator of the global cell metabolism. We then explore the likely consequences of this dependency on the evolution both of cytosine-related innate immunity processes and of the viral genome sequence. Subsequently, we delineate critical details of the impact on the virus biological functions on the nucleotide composition of its genes and consequences for its evolution.

## RESULTS

### In-depth analysis of pyrimidine metabolism highlights the unique position of CTP in metabolism

To understand how viruses recruit the functions of their host cells to their benefit, we must understand what would be the point of view of a virus if it were to sustain continuous propagation. Essentially lacking biosynthetic potential, a virus must tap into the host’s metabolic resources. This introduces a considerable limitation in the metabolic options offered to viral replication. For this reason many viruses ended up coding for functions that are missing or deficient in their hosts (9). Some even help their hosts to upgrade their built-in ability to make the most of their environment, thus ensuring a wealthy propagation of the viral progeny. Auxiliary metabolic genes are commonplace in bacteriophages (10), but also in a variety of eukaryotic viruses, such as herpes viruses (11). Selection pressure *via* efficacy of transmission multiplied by number of replicates per cell, coupled to selection stemming from intracellular availability of essential precursors (nucleotides, amino acids, lipids and carbohydrates) creates a variety of bottlenecks that shape the virus evolutionary landscape (12–14). Furthermore, the envelope of many animal viruses is built up from components of the host cell’s membranes (15, 16), as well as a capsid made of virus-specific proteins (17, 18). To harden them against environmental offences and provide them with addressing tags, some of these proteins are glycosylated, which involves tapping into the cell’s resources of UDP-sugars and GDP-sugar precursors (19, 20). Besides this uracil metabolism-dependent protective and tagging feature, we highlighted here the relevance of the membrane lipids, which uniquely derive from precursors involving pyrimidines, specifically liponucleotides based on a CDP skeleton (21–23). We also emphasized the need for a virus to express its protein to have an active tRNA complement, necessitating the function of host CCAse. How the construction of the building blocks are put together in an uninfected cell is therefore the very first challenge faced by the virus after it has accessed the cytoplasm of the host cell. Finding an answer to this question dictates the exploration of the mystery of the cells’ assembly lines that prepare them for growth. Before exploring quantitative consequences of this unique design, we summarize how cytosine-based metabolism is organized in the next couple of paragraphs.

#### The logic of energy management for nucleic acids synthesis

Synthesis of the viral RNA genome draws resources not only from the metabolism of pyrimidines but also from the general logic of the cell’s energy management. A key chemical feature of the related processes is that they rest on hydrolysis or synthesis of phosphate bonds (24). Versatility of these processes is ensured by the usage of the shortest polyphosphate structure, that of nucleoside triphosphates, NTPs—we do not consider here the special case of purely mineral polyphosphates (25). General metabolism is developed along two lines with respect to its energy demands (this is illustrated in **Figure 1** for pyrimidine metabolism).

**Figure 1.**
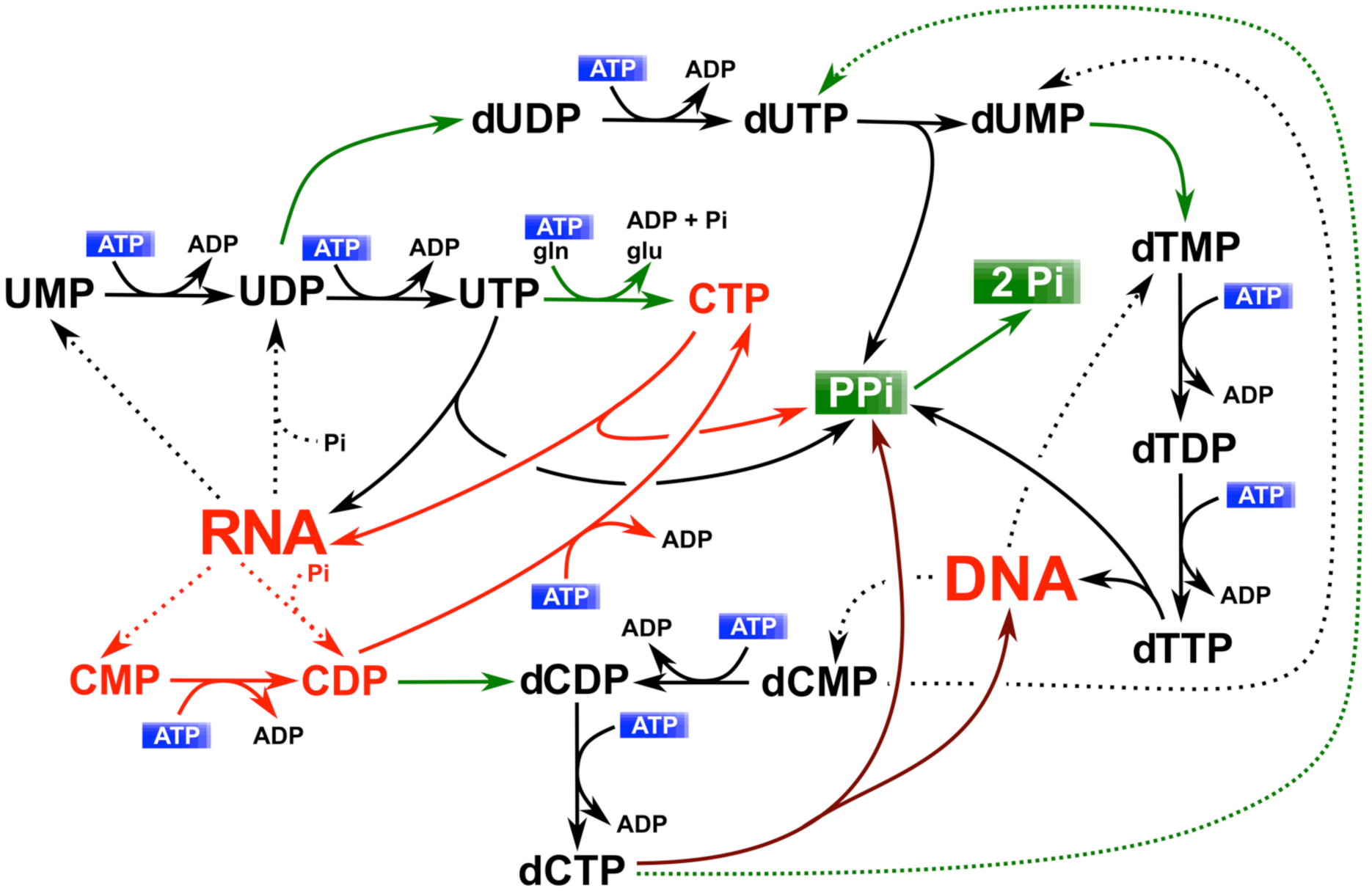
Energy-driven pyrimidine-based nucleic acid metabolism. ATP is the general donor in the biosynthesis of pyrimidines. CDP, required as a precursor of dCTP synthesis is produced by RNA turnover via hydrolysis or phosphorolysis (red arrows). RNA and DNA synthesis is driven by pyrophosphate hydrolysis (green arrows indicate irreversible reactions). dTTP is results from a pyrophosphate-driven reaction producing dUMP, and is finely tuned by thymidylate kinase, which makes its immediate precursor dTDP.

When energy is meant to be used in a reversible way, NTPs are hydrolysed into NDP+Pi. This is where the role of mitochondria is critical in non-proliferating cells (26). These organelles restore the ATP complement of the cell, in particular to the endoplasmic reticulum—ER, (27), and in the present context this is crucial for the generation of new viral particles. In contrast— and this is relevant not only for intermediary metabolism but also for macromolecule biosynthesis, with more than 500 such reactions reported in the KEGG database (28)—when the relevant pathways have to be driven forward, triphosphate hydrolysis produces pyrophosphate (PPi). PPi is subsequently hydrolysed irreversibly into two phosphates by omnipresent pyrophosphatases: NTP => NMP + PPi => NMP + 2 Pi, and this drives syntheses forward. Biosynthesis of macromolecules rests to a great extent on this two-pronged strategy.

Diverging from this general design, DNA synthesis is an exception in most organisms. In this case, dNTP synthesis is indirect. It is a two steps process. NDPs are used instead of NTPs as precursors of the dNDPs, which are subsequently phosphorylated to dNTPs (29). The underlying metabolic logic is that, if DNA synthesis used NTPs instead of NDPs, the nucleotide availability would build up a considerable metabolic pressure driven by the cellular concentration of nucleotide precursors. This would affect genomes by expanding progressively their length —which already usually occupies a sizeable portion of the cell’s volume —from generation to generation. Using NDPs alleviates the difficulty: nucleotide reductases evolved so as to use ribonucleoside *di*phosphates not *tri*phosphates to make deoxyribonucleotides (30–32). The concentration of NDPs is of the order of 1% that of NTPs because metabolism is poised to supply energy via steady synthesis of ATP, used to keep the other NTPs at a high level as well. This lowers by two orders of magnitude the metabolic pressure on DNA synthesis. Yet, using NDPs in this pathway leads to a logistic metabolic cost, and this is specific to cytosine nucleotides, placing them in a singular position in metabolism. *De novo* synthesis of cytidine triphosphate (and hence of all cytosine derivatives) does not go via a CDP intermediate. It is the direct product of amidation of UTP (33), using glutamine as the nitrogen donor and ATP, with CTP synthetase as a catalyst [EC 6.3.4.2 (34)]. This functional arrangement requires specialized biochemical reactions in order to maintain a significant level of cytosine in DNA sequences. Synthesis of CMP (and subsequently, or in parallel, CDP) is only achieved via RNA and membrane lipid precursors turnover (red arrows in **Figure 1** and red and purple arrows in **Figure 2**). A widespread consequence of this feature is that DNA-based genomes tend to become progressively more AT-rich—note that depleting C in double stranded DNA genomes must be followed by G depletion because of the first Chargaff’s parity rule (35)—in a variety of conditions (36).

**Figure 2.**
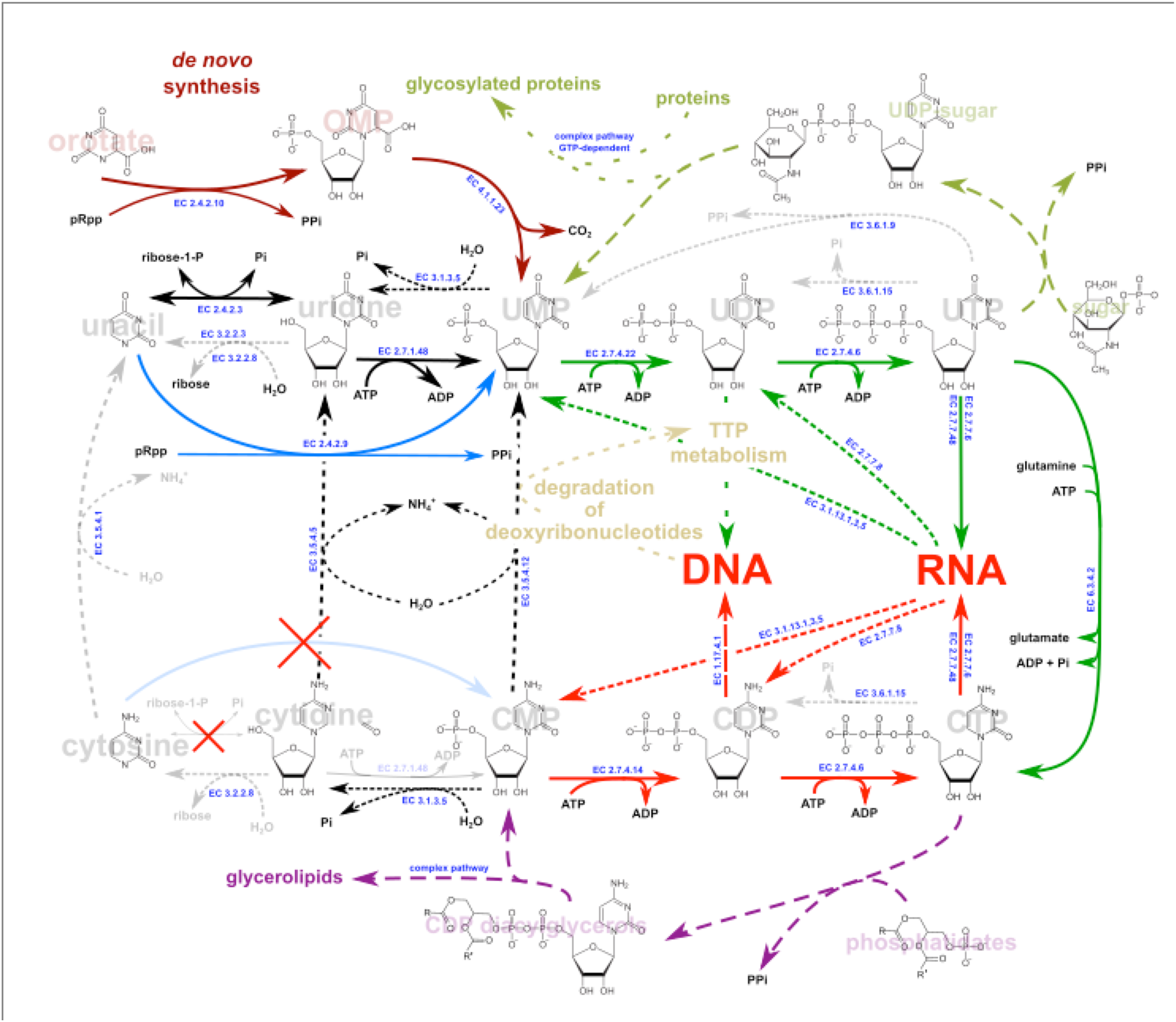
Synthesis and salvage of pyrimidine nucleotides. Synthesis of UMP begins with orotate phosphoribosyltransferase followed by decarboxylation (brown arrows). The anabolic pathway ends up with UTP and CTP (green arrows). Salvage of CTP stems from RNA metabolism (red arrows) and lipid metabolism (purple arrows). Degradation and scavenging of cytosinebased nucleotides goes through uracil-based scavenging and return to the CTP biosynthetic pathway, with cytosine deamination of intermediates as critical steps. Uracil phosphoribosyltransferase matches the role of the orotate counterpart in the ultimate salvage of the base (blue arrow). No counterpart has been yet identified, to our knowledge (see text), for scavenging cytosine (crossed out light blue arrow). Distribution of relevant enzymes in H. sapiens and model bacteria is illustrated in Table 1.

#### An unexpected secret of life: cytosine metabolism provides both a rheostat and a flywheel to integrate growth of the various cell structures, constraining coronavirus development

Does this original role of cytosine nucleotides extend to RNA genomes? Not in any straightforward way: synthesis of RNA utilizes NTPs, not NDPs. The singular requirement for NDP precursors for DNA synthesis might even be a selective force that drove the evolution of viruses away from DNA-dependent forms, despite the high intrinsic instability of RNA as compared to DNA. The dCDP bottleneck would prevent DNA viruses to reach a high virus burst size and could make easier the emergence of some kind of innate immunity. It would also favour emergence of viruses based on RNA metabolism and this should be taken into account when exploring the origin of viruses (37, 38). Yet, we may ask: are RNA viruses still submitted to patent metabolic constraints, and what would they be? At first sight, the answer looks negative, at least involving nucleotide biosynthesis, because *de novo* synthesis of nucleotides allows direct production of each of the nucleoside triphosphates. All the same, there might remain some persisting negative trend against cytosine accumulation, because CTP synthesis requires both ATP and a nitrogen source, which makes availability of the molecule highly sensitive to energy and nitrogen availability (**Figure 2**). However, this type of negative pressure equally applies to synthesis of ATP, for example (39). Then, if anabolic processes are not preset to modulate the stoichiometry of specific NTPs, what about their degradation and salvage?

**Table 1.**
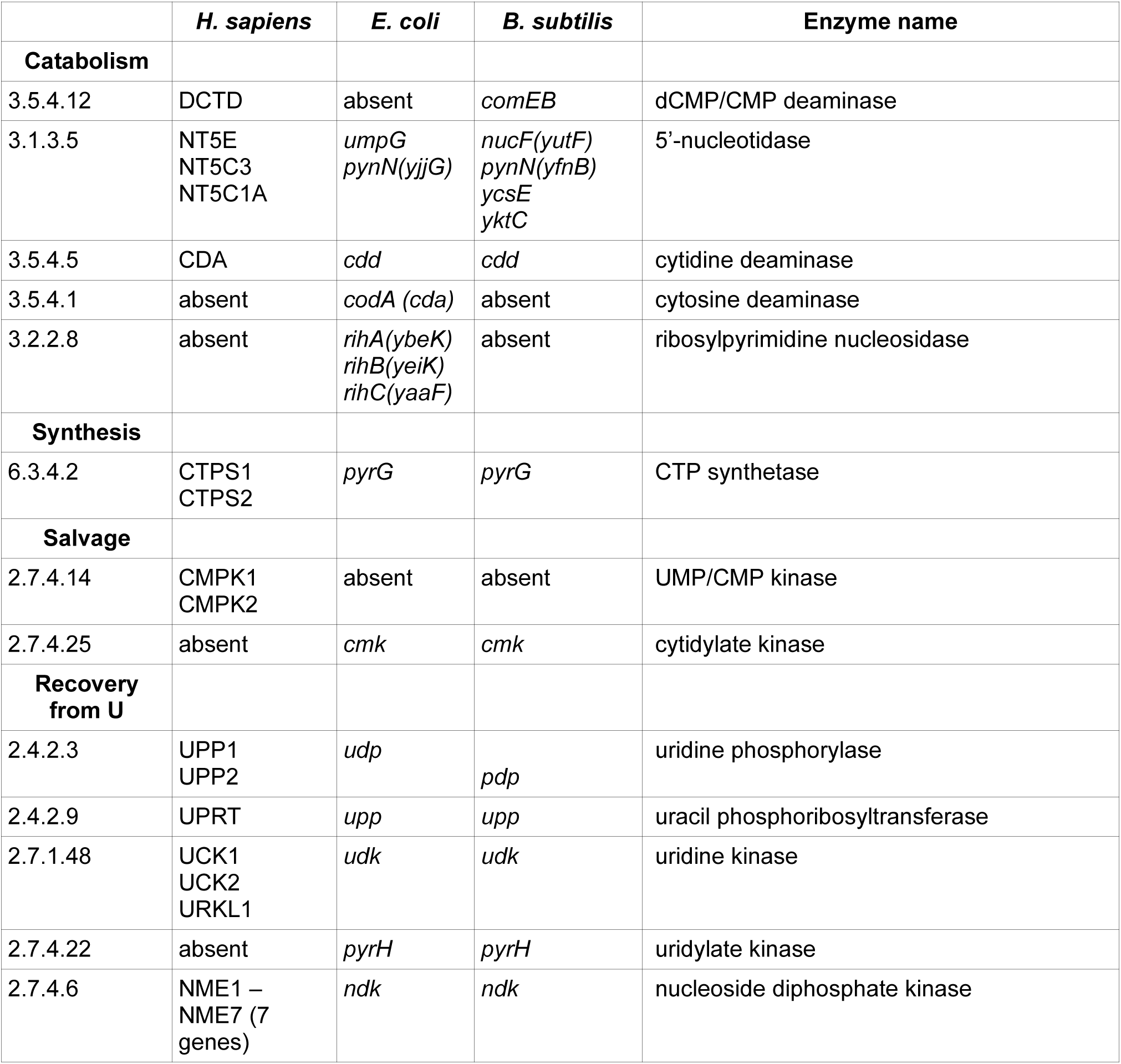
Cytosine-related salvage in human cells: present and missing enzymes with bacteria for comparison

All metabolites wear off with time. This implies that they must be degraded and recycled, either as a whole or as parts. In the case of ribonucleotides, three units—a phosphate, a ribose, and a heterocyclic base—can go through specific degradation or salvage pathways. During DNA turnover or following proofreading steps, a fast track converts CMP to CDP then CTP, but in the case of dCMP, while this track exists, there is a parallel, unexpected one that begins with deaminating dCMP into dUMP, which subsequently feeds into the TTP biosynthetic pathway—see (40, 41) and **Figure 1**. Another pathway produces cytidine after hydrolysis of the 5’-phosphate of CMP, then deaminates it to uridine (42), or, in bacteria but not in multicellular organisms, producing cytosine then deaminated to uracil (43), which is mainly scavenged directly by uracil phosphoribosyltransferase (EC 2.4.2.9, **Figure 2** blue arrow) directly into UMP. That this indirect route plays a crucial role in cells is witnessed for example in the fact that the whole RNA-derived salvage pathway (cytidylate phosphatase and cytidine deaminase) is critical in embryonic development (44). This is because any imbalance in dNTP pools is highly mutagenic (45). Furthermore, the enzymes of this salvage pathway are also important to recycle the modified derivatives of cytosine which result from frequent metabolic accidents or are encountered as epigenetic markers (46).

Remarkably, the salvage pathways are straightforward for all nucleobases, cytosine excepted (**Table 1**).

The purine salvage pathways have been thoroughly explored, in particular in animal pathogens and in plants (47–49), while the pyrimidine salvage pathways have remained somewhat less studied (50). The omnipresent roles of ATP and S-adenosylmethionine (AdoMet) often ends up with adenine as a waste product. As a consequence, natural selection retained a variety of enzymes meant to scavenge adenine wastes, so that any downwards trend in the ever critical ATP supply would be easily overcome—see *e*.*g*. (51, 52). A key enzyme in this salvage process is adenine phosphoribosyltransferase [EC 2.4.2.7, (53), 1286 references at PubMed on 18/06/20]. In the same way, guanine or uracil are salvaged *via* (hypoxanthine)-guanine phosphoribosyltransferase [EC 2.4.2.8, 1536 references at PubMed on 18/06/20, (54)], or uracil phosphoribosyltransferase EC 2.4.2.9, (55), 310 references at PubMed on 18/06/20]. Further comparable activities exist, such as orotate phosphoribosyltransferase EC 2.4.2.10 (56)—which inputs orotate at the beginning of the pyrimidine biosynthetic pathway—and the somewhat less similar nicotinamide phosphoribosyltransferase EC 2.4.2.12 (57). These enzymes share a common descent, showing that life has easily evolved a panoply of related activities.

It was therefore expected that the same would hold true for cytosine, allowing the cell to scavenge the base easily from its environment. Yet, it appears that no cytosine phosphoribosyltransferase exists in any extant organism (light blue arrow, **Figure 2**). Preliminary work with the protozoon *Giardia lamblia* suggested that this activity might be present in the organism (58, 59). Surprisingly however, deciphering the genome strongly suggested that CTP was derived from cytosine deamination into uracil, followed by salvage of uracil and amidation. Indeed in the most recent release of GiardiaDB, there appears to be no sequence in the genome of the organism that could be attributed to cytosine phosphoribosyltransferase—see (60) for access to the genome database. At this point, therefore, no extant organism codes for such an enzyme. By contrast, despite the lack of *de novo* biosynthesis pathways for pyrimidines in this organism, the genome still codes for a CTP synthetase (PyrG). This is in line with the general observation that cytosine and related cytosine-containing derivatives are systematically deaminated into uracil-containing derivatives, subsequently processed to regenerate CTP (**Figure 2**, and **Figure 1** for salvage of processed DNA derivatives). As a further case in point *pyrG* has also been found as a necessary complement required for life in the smallest genome of an autonomous synthetic construct (61). This strongly suggests that recovering CTP requires a uracil-dependent pathway as well as that independent management of cytosine-based nucleotides is critical to govern metabolism, even in the presence of a rich supply of metabolites from the outside—note that *Giardia* is a parasite.

This singular positioning of CTP synthesis in metabolism makes CTP synthetase a convenient enzyme to adjust the flow of cytosine-containing nucleotides, acting as a rheostat does in an electric contraption—see *e*.*g*. (62). Substantiating this unique role, the functional structure of the enzyme displays a very unusual architecture. It makes filaments, named cytoophidia—specific membrane-less organelles that control the spatial distribution of cytosine-dependent intermediary metabolism (63, 64)—in all the organisms where its organization has been explored (65). The structure of CTP synthetase is important in the present context because the synthesis of membrane lipids is a further metabolic step that involves the nucleotide, with most membranes deriving from cytosine-based liponucleotides (66, 67). While the precise membrane lipid composition differs in the three domains of life, the general organisation of the cognate pathways is similar. Because the lipid content of cells can vary over a wide range, the stores of CDP-containing liponucleotides, in particular in eukaryotes—with an important network of intracellular membranes—is preset to play the role of a flywheel, allowing fine tuning of the availability of cytosine-derived metabolites in the cell when conditions vary. This property could have been advantageously recruited for innate immunity. Finally, as perhaps could be expected at this point of our demonstration, the very first enzymes for *de novo* synthesis of pyrimidines, carbamoyl-phosphate synthetase (*C*PSase), aspartate transcarbamylase (*A*TCase), and dihydroorotase (*D*HOase) are associated into a multifunctional structure, named CAD (68). Besides a general role in management of cell growth, CAD is highly expressed in leukocytes, where it enables Toll-like receptor 8 expression in response to cytidine and single stranded RNA (69), a situation met upon infection by RNA viruses. Supporting a role of CAD in antiviral innate immunity, its activity is modulated by a dedicated viral protein during Enteroviral infection (70).

### Consequences of cytosine-related imbalance in the nucleotids composition, evolution and coding capacity of coronavirus genomes

The most straightforward consequence of the metabolic qualitative design just outlined is that a metabolic force will keep driving the cytosine content of RNAs to lower values, unless opposite processes—and selection pressure leading to discard organisms with too low cytosine content, for example because this would create unbearable biases in the amino acid composition of the proteins coded by these genomes— had the upper hand during evolution. This prompted us to develop a brief quantitative analysis of pyrimidine metabolism in relation with SARS-CoV-2 infection, as we now document.

#### Cytosine content-related phylogeny of some virus isolates

The constraint on cytosine availability witnessed in the composition of coronaviruses—in particular SARS-CoV-2—is likely to reflect the coupling between synthesis of viral particles and the host cell’s metabolic capacity. In order to assess the evolution of the virus with these metabolic constraints we decided to evaluate the C content using 89 representative strains from the four genera of coronaviruses, which were selected based on their phylogenetic and host background. Here we developed two distinct approaches in order to take into account 1/ the nucleotide patterns across the virus group, and 2/ the coding sequence-related limitations that constrain the function of the viral proteins as they adapt to their host.

To study the evolution of coronaviruses, we used standard techniques to generate a phylogenetic tree of representative strains (see **Methods** section), where we highlighted the average cytosine content of each viral genome (**Supplementary Figure 1**). There is significant variation between the cytosine content of viruses of different clades. Based on 89 representative genomes we found that coronaviruses have 27.3% A, 17.9% C, 21.5% G and 33.3% U on average. Overall, the coronavirus genomes contain a bit more pyrimidines than purines, with a mean content for C+U of approximately 51.2%.

Currently, seven coronaviruses are known to infect humans. These include four epidemic CoVs causing mild respiratory symptoms in humans (HCoV-229E, HCoV-NL63, HCoV-HKU1 and HCoV-OC43), the severe acute respiratory syndrome coronavirus (SARS-CoV-1) causing the pneumonia outbreak during 2002-2003 in China (71), the Middle East respiratory syndrome coronavirus (MERS-CoV) still causing small outbreaks through camel-to-human transmissions in the Middle East and the newly emerged SARS-CoV-2 (72). These viruses all originated from bat species, although the exact intermediate hosts involved in human infections remains obscure (73–77). Interestingly, the cytosine content of bat viruses is variable (from 16.0% to 21.5%), reflecting the diverse virus species harboured by bats, which make a very diverse natural reservoir for many coronaviruses. Intriguingly, we observed a lower cytosine content for the four human established epidemic viruses (HCoV-HKU1, 13.0%; HCoV-NL63, 14.4%; HCoV-OC43, 15.2%; HCoV-229E, 16.7%) than the three spillover-related viruses infecting humans (SARS-CoV, 20.0%; MERS-CoV, 20.3%; SARS-CoV-2, 18.4%) and most bat viruses. The four human epidemic viruses have been established in human populations for decades. The significantly low cytosine content observed for these two viruses may indicate host-related features impacting cytosine-containing metabolites availability. Several studies of the codon usage biases in emerging coronaviruses have been published, providing a general view of the situation—see e.g. (78), without, however, exploring the contextual constraints in terms of metabolic properties of their hosts. If the adaptation of coronaviruses to the human species was to drive down the cytosine level in the virus genome, then we might expect the cytosine content of SARS-CoV-2 strains to decrease as the epidemic unfolds. Our regression model is consistent with this view (**Figure 3**).

**Figure 3:**
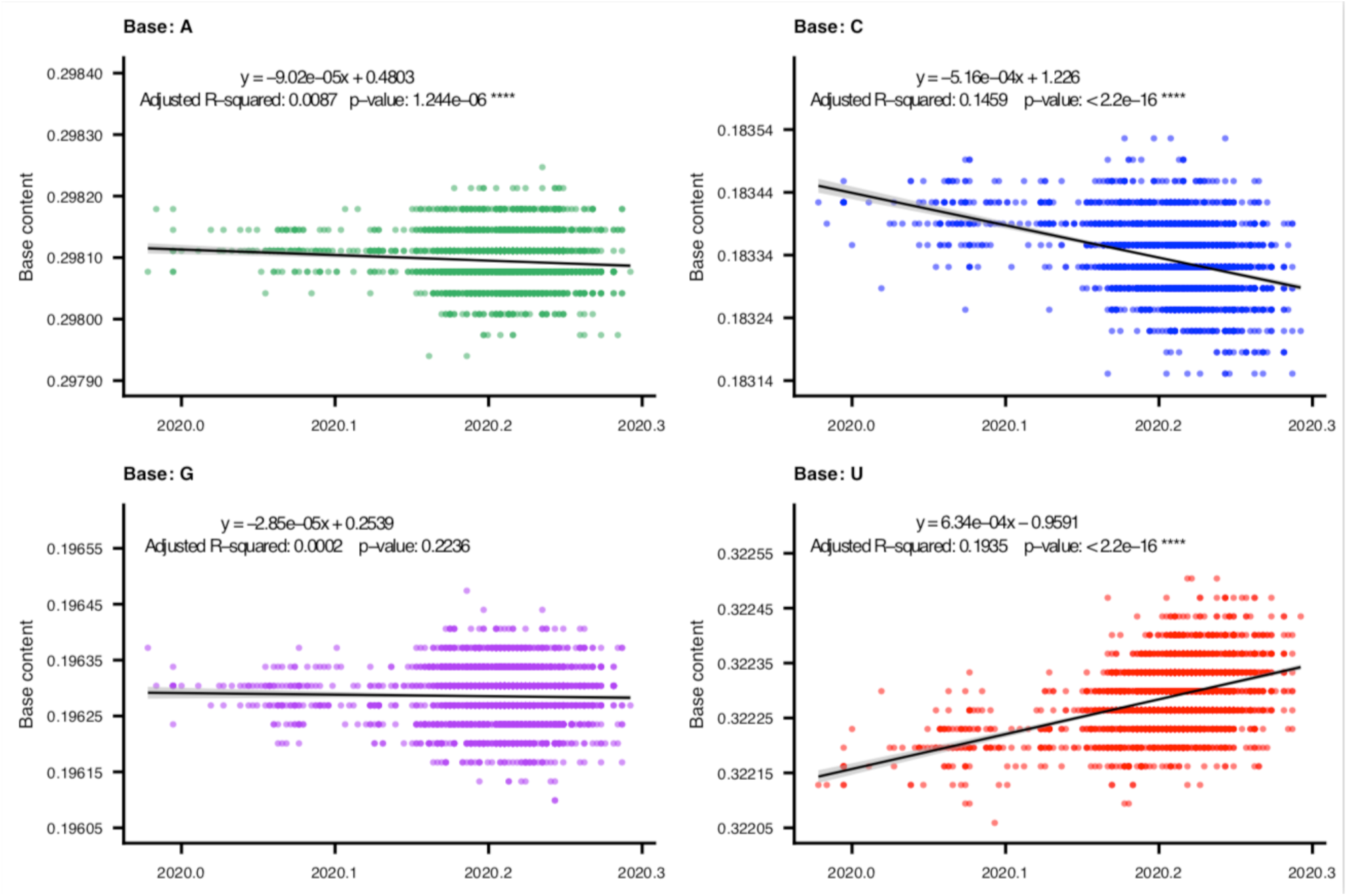
Dynamic of base composition at the coding regions of SARS-CoV-2. The base composition of the coding regions concatenated by 26 ORFs of SARS-CoV-2 is displayed. Each dot represents one sequence. The calculation was based on 2,574 unique SARS-CoV-2 strains isolated from December 24^th^, 2010 to April 17^th^, 2020.

It allowed us to estimate that SARS-CoV-2 may lose its C complement by 0.000516 base per position per year (y = −0.000516x + 1.226, adjusted R^2^ = 0.1459) while gaining U by 0.000634 per year (y = 0.000634x - 0.9591, adjusted R^2^ = 0.1835), under its current circulation dynamics in susceptible populations. A parallel increased trend for A and a decrease for G was also observed, but with more moderate slopes than those for C and U. A Wilcoxon rank sum test showed that the content of the four bases in the SARS-CoV-2 sequences was significantly different in each case, while somewhat correlated with each other. The strongest correlations were observed between two groups (**Supplementary Figure 2**): a decrease in A was significantly correlated with an increase in G (adjusted R^2^ = 0.5738), while a decrease in C was significantly correlated with an increase in U (adjusted R^2^ = 0.7177). As a consequence the SARS-CoV-2 virus is on its way to gradually lose C during its adaptation in humans, resulting in a genomic base composition more like those of the four previously established human endemic coronaviruses.

The second approach did not aim at creating a phylogeny of the viruses, but, rather, a cladistic tree showing structural properties shared by viruses that are likely to indicate affinity of functional features. This approach assumes that, after sufficient time of evolution, the nucleotides present at the majority of sites in the sequence have had chances to be modified several times reaching local saturation, so that it is difficult or impossible to link those with specific features of the proteins encoded in the virus (the beginning and end of the sequence, that are critical for RNA-dependent replication are not taken into account). By contrast, the presence of insertions or deletions (indels) will affect considerably the overall structure of the proteins, and this should impact their function in a way that is not likely to be reversible—see for example (79). An earlier such report (80) demonstrated the usefulness of this approach.

The use of cladograms in this case attempts to show the relative distances and should not necessarily reflect the evolutionary history of the group. Compared to the standard alignment of the 89 reference genomes (**Supplementary Table S1** and **Supplementary Figure 1**), the equivalent gap-based alignment uses undefined characters for nucleotides and ‘dummy’ characters for gaps to cheat the algorithms for tree construction so that distances are calculated on the basis of the sums of scores for gap positions (presence of indels, see **Methods**). While it is not possible to check one by one each and every gap, the common ones are likely to reflect a common structure or function characteristic of the corresponding region (**Figure 4**).

**Figure 4.**
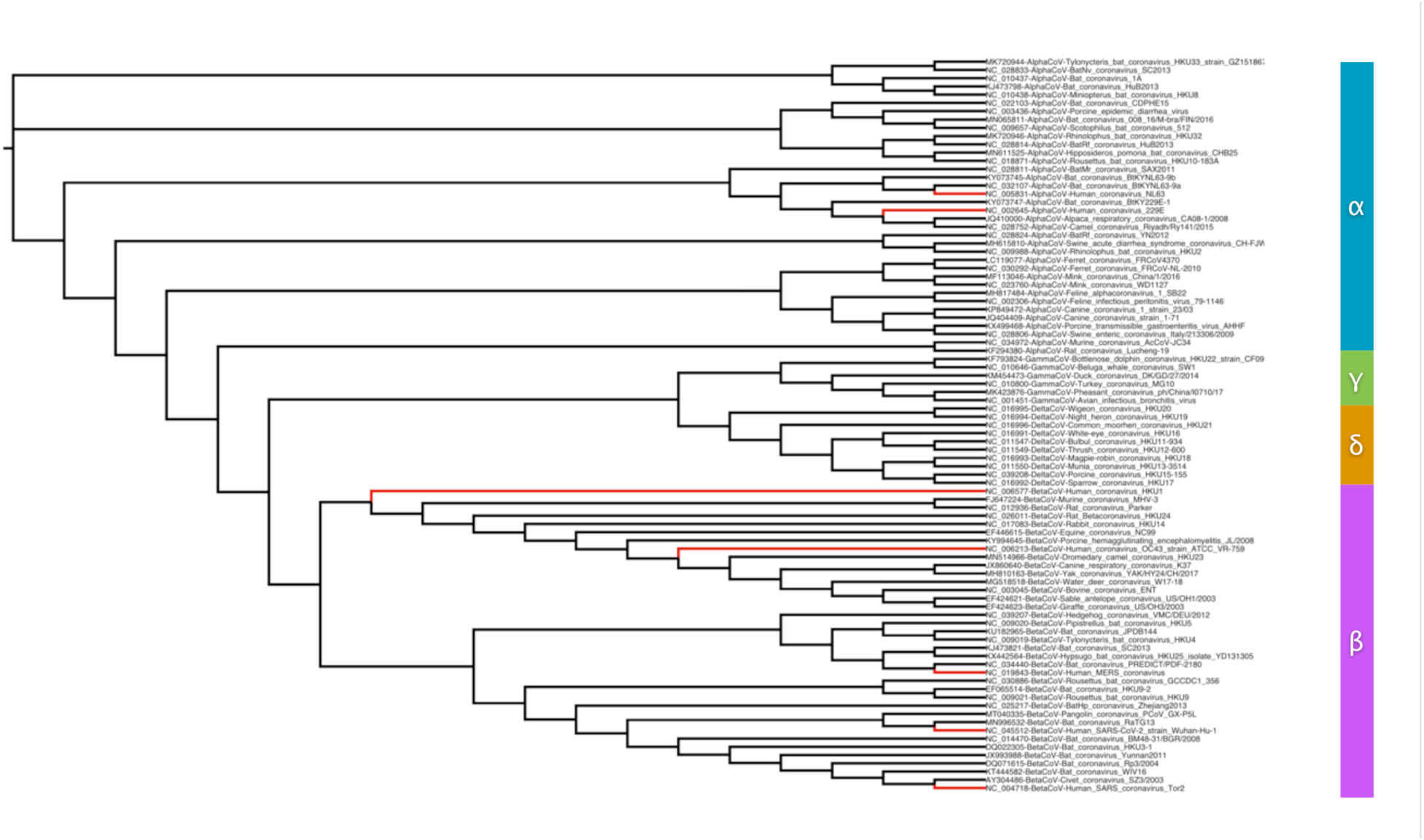
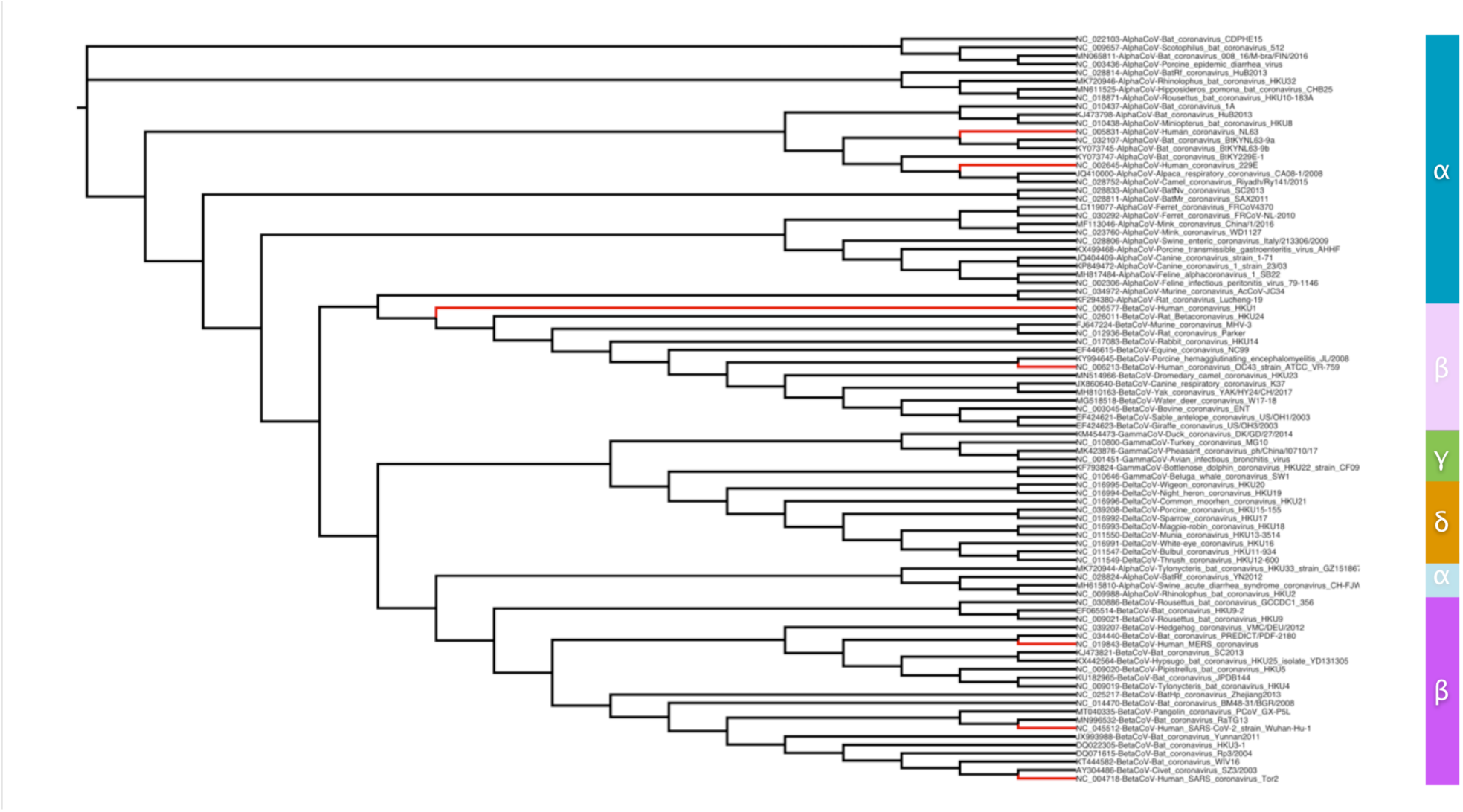
Phylogeny of coronavirus representatives based on full genome sequences (panel A) and on indels (panel B) A genome-based tree is generated based on the full genome sequence alignment of the group (panel A) and a gap-based tree is created, based on insertions and deletions (indels) only (see Methods). The seven known human coronavirus strains are highlighted by a red colour for the corresponding branches. A panel on the right indicates the four coronavirus groups; in panel B, the two incongruent sub-groups are shown by the same colour code with reduced opacity (alpha and beta sub-groups). For details, please see text.

Remarkably, both the standard, genome-based tree and the gap-based tree are quite congruent—i.e. knowing only the indel content is enough to draw a tree that describes the relationships between the coronavirus groups (**Figure 4**). In the genome-based tree, the four groups of coronaviruses are detected clearly, and the seven human virus strains are highlighted, in groups alpha and beta (**Figure 4a**). In the gap/indel-based tree, the four groups are also consistently derived, with the exception of a beta sub-group that contains the two human viruses with reduced pathogenicity potential, compared to the beta sub-group that contains the SARS, MERS and SARS-2 strains (**Figure 4b**). At the same time, a tiny alpha sub-group exhibits similar indel patterns with the latter beta sub-group, presumably with similar indel patterns. It is tempting to speculate that these alpha sub-group strains may share certain hitherto unknown properties that might render them potentially dangerous in terms of zoonotic disease capacity. The observed patterns could be interpreted as showing that what is coded in the indel regions has a considerable weight on the virus adaptation to their hosts and does not strictly depend on the base composition and amino acid coding potential of the genome sequences. More research is needed to establish the nature of indel-based trees in the future.

#### Biased codon composition of the regions coding for individual viral proteins

Coronaviruses, and many positive-sense single-stranded RNA viruses as well, produce plus strands at a 50- to 100-fold excess of their minus-strand replicated template. Because several regions in the 3’ half of the virus are « transcribed » from the template RNA minus-strand of the virus (81), a further deviation from parity should appear in the nucleotide usage for virus construction. This means that the overall nucleotide consumption is not strictly constrained by the second Chargaff’s parity rule, that would result in an amount of A equal to that of U, and G to that of C (82). The virus multiplication rests on a RNA-dependent replication process, so that any pressure on a given base availability—here C—would affect its complement—G in our case. As discussed in the previous section, we expected a general selection pressure operating on CTP and tending, in the long run, to decrease the C content of the RNA virus, but also that of G.

Furthermore, this implies a particular imbalance in the nucleotide composition of the viral RNA, allowing it to differ from standard mRNAs of the host cell. We therefore expect that the virus will interfere with the host’s translation machinery in a way that allows it to be discriminated positively against the cell’s mRNAs (see **Discussion**). This should have consequences for the translation of the viral genome. The virus encodes proteins that have essential functions for its development, not only for the replication machinery and the formation of a capsid, but also for several ancillary functions needed for hijacking the host metabolism. The constraint on the genome nucleotide composition must be reflected in the codon usage bias of the virus protein coding sequences, with important consequences on the way tRNAs are used. Furthermore because natural selection acts on viral functions, it can be expected that the outcome of the general C lowering trend will differ in different proteins encoded by the virus, depending on the selection pressure constraining their functions. Taking advantage of the degeneracy of the genetic code, SARS-CoV-2 could also limit its C content *via* the use of alternatives at the third codon position by selective codon usage.

To explore this hypothesis, the relative synonymous codon usage (RSCU) values were calculated for each coding region of SARS-CoV-2 to reveal any differential usage of synonymous codons. An RSCU value of 0, 0∼0.6, 0.6-1.6 or >1.6 implies that a codon is not-used, under-represented, normally-used, or over-represented—respectively for the four value ranges (83). Among the 26 major coding regions, six (Nsp11, ORF10, ORF7b, ORF6, E and Nsp7) have a translated peptide length of less than 100 amino acids, which would result in codon usage profiles of low statistical significance. For example, Nsp11 encodes only 33 amino acids, which explains the very limited number of codons utilized by the corresponding gene. These sequences were not further analysed (**Figure 5**). The three codon positions do not have the same importance in the selection of specific amino acids. Pressure against C must have consequences in terms of the protein coding landscape of the virus. In general, in regions where tRNA choice allows a wobble between U and C, the virus sequence is considerably enriched in U. It seems noteworthy that the codon usage bias and bias in tRNA choice differs between genes of the human host and the virus genes, except for two viral proteins, accessory protein Nsp1 and nucleocapsid protein N, notwithstanding preference of U over C—e.g. preference for GGU and CGU codons in protein Nsp1 (**Supplementary Figure 3**).

**Figure 5.**
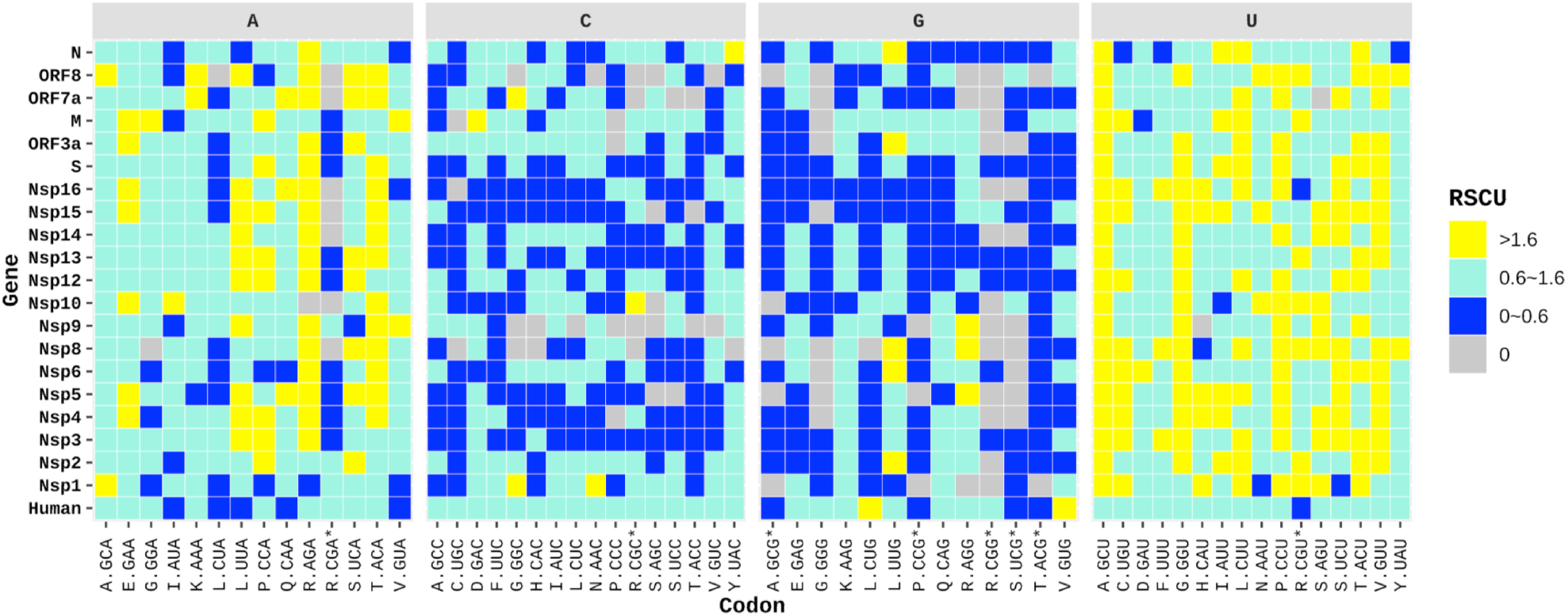
Codon usage of SARS-COV-2 ORFs based on the third codon position compared to human coding regions. The 20 SARS-CoV-2 ORFs with a length of over 300 nucleotides were submitted to RSCU calculation. The codon usage for 120,426 human coding regions was also displayed to facilitate comparison. Codons are displayed with the first letter denoting the amino acid and the three letters following a dot representing the codon. Codons containing a CpG dinucleotide are suffixed by an asterisk. The four panels separate the codons according to the nucleotide located at the third codon position. Codons that are not used (RSCU=0), under-represented (RSCU<0.6), normally utilized (RSCU ranges between 0.6-1.6), or over-represented (RSCU>1.6) are labeled in gray, blue, ice cold and yellow, respectively.

The first C codon position is used to input histidine, glutamine, proline, and arginine or leucine in proteins. This is particularly significant for the proline residue, essential in the folding of key viral protein domains (Li *et al*., 2014), because it is encoded by CCN codons. All proteins of SARS-CoV-2 prefer the usage of CCA or CCU codons for proline, avoiding the usage of C and G (**Figure 5**). Histidine and glutamine are in two-codon boxes, discussed below. Arginine presents a different situation because CGN codons can be replaced by AGR codons: SARS-CoV-2 favours the AGA codon especially, with only one compulsory G. AGG codons are enriched in proteins Nsp5, Nsp8 and Nsp9. Remarkably however, the Nsp1 protein, which corresponds to the initial domain of the ORF1a(b) protein, and is translated very early on in the virus expression cycle, contains only CGH codons (H = A, U or C) and this is in total contrast with the other viral proteins (except for Nsp10, yet this protein has only two arginine residues, making this observation possibly irrelevant). Codon CGG is only present 11 times in the coding sequences of the virus, suggesting that when present, it has been submitted to positive selection, possibly at a site important for the translation-coupled folding of the protein. The most interesting location of this codon is a doublet that corresponds to a four codon insertion in the spike protein of the virus. Finally, the pressure on leucine content is also lower. CUN codons are used to code for leucine, with the majority using codon CUU, but this amino acid can be introduced using the alternative UUR codons. Yet, UUA is used more frequently than UUG. UUG is relatively enriched in proteins Nsp6, Nsp8, ORF3a and N (**Figure 5** and **Supplementary Figure 3**).

In the second position requiring a C we find proline again, and also threonine (ACN), alanine (GCN) and serine (UCN). For threonine, codon ACU is the most used codon, progressively being generally replaced by ACA as we progress along the genome sequence, ACC and ACG are rare. Alanine is mainly encoded by GCU codons, followed by GCA, with protein Nsp1, again, differing somewhat from the other viral proteins in that the frequency of GCA and GCU are the same. Serine (UCN) is able to escape much of the constraint imposed by C availability as it can use the alternative AGY codons. Codons UCU and UCA are more or less used in an equivalent way, except, again, in protein Nsp1, which mainly uses AGU and AGC codons. In general AGC is seldom used while AGU is the dominant serine codon.

Finally, the third position can be replaced by A, U or G in the four codon boxes, two of which, valine and glycine, are further discussed below. In general the corresponding NNC codons are rarely used. Again, protein Nsp1 is an exception, with codon GGC used more frequently than GGU. Overall GGU is dominating with some contribution of GGA, while GGG is often absent. In the case of valine (GUN codons), the dominating codon is GUU, followed by GUA. By contrast, U-ending codons which are rarer than expected are clustered in several proteins: UGU, UUU and UAU in protein N; AUU in Nsp10; CGU in Nsp16; GAU in protein M, CAU in ORF8 and AAU and UCU in Nsp1. NAU codons correspond to two codon boxes (NAN codons). These codons are discriminated along a pyrimidine / purine axis. A pyrimidine (NAY) is used to maintain the same nature of the coded residue whether the codon uses a U or a C as its 3’ end (aspartate, asparagine, histidine and tyrosine), while a purine (NAR) allows coding for glutamate, glutamine and lysine. UAR codons are also specifying the terminal step of translation. As stated above, the SARS-CoV-2 genes avoid the usage of C containing codons whenever possible (**Figure 5**). Moreover, probably due to the base-pairing requirement imposed during transcription and replication, the virus also avoids the usage of G-ending codons. This avoidance is maintained in the overall choice of NAR codons, except in protein Nsp6 where CAG is preferred to CAA, as well as GAG over GAA and CAA over CAG, which suggests that this results from a significant selection pressure. This is the more remarkable because Nsp proteins are cleaved off large ORF1a and ORF1ab precursors. In general, and this is as expected, codons NAU are preferred over NAC for the pyrimidine ending codons of NAN boxes. The exceptions are, for GAC, protein Nsp5 and protein M; for CAC, Nsp10 and ORF7a; for AAC, protein Nsp1 and protein M; and for UAC, proteins Nsp1, Nsp5, Nsp9, M, Orf7a and N.

#### tRNA-dependent modulation of synonymous codon translation

Consistent with metabolic pressure against CTP and despite their uneven coding length, most of the genes of SARS-CoV-2 avoid the usage of the C-ending codons (**Figure 5** and **Supplementary Figure 3**). A similar low preference for C-ending codons was also observed for coronaviruses of other species and genera, as calculated using the ORF1ab coding region (**Supplementary Figure 4**). The way tRNAs are utilized as a function of the first anticodon (third codon) nucleotide is highly unsymmetrical. For this reason the corresponding tRNA position (N34) is usually heavily modified, while specific tRNAs are deciphering individual codons (**Table 2**).

**Table 2.**
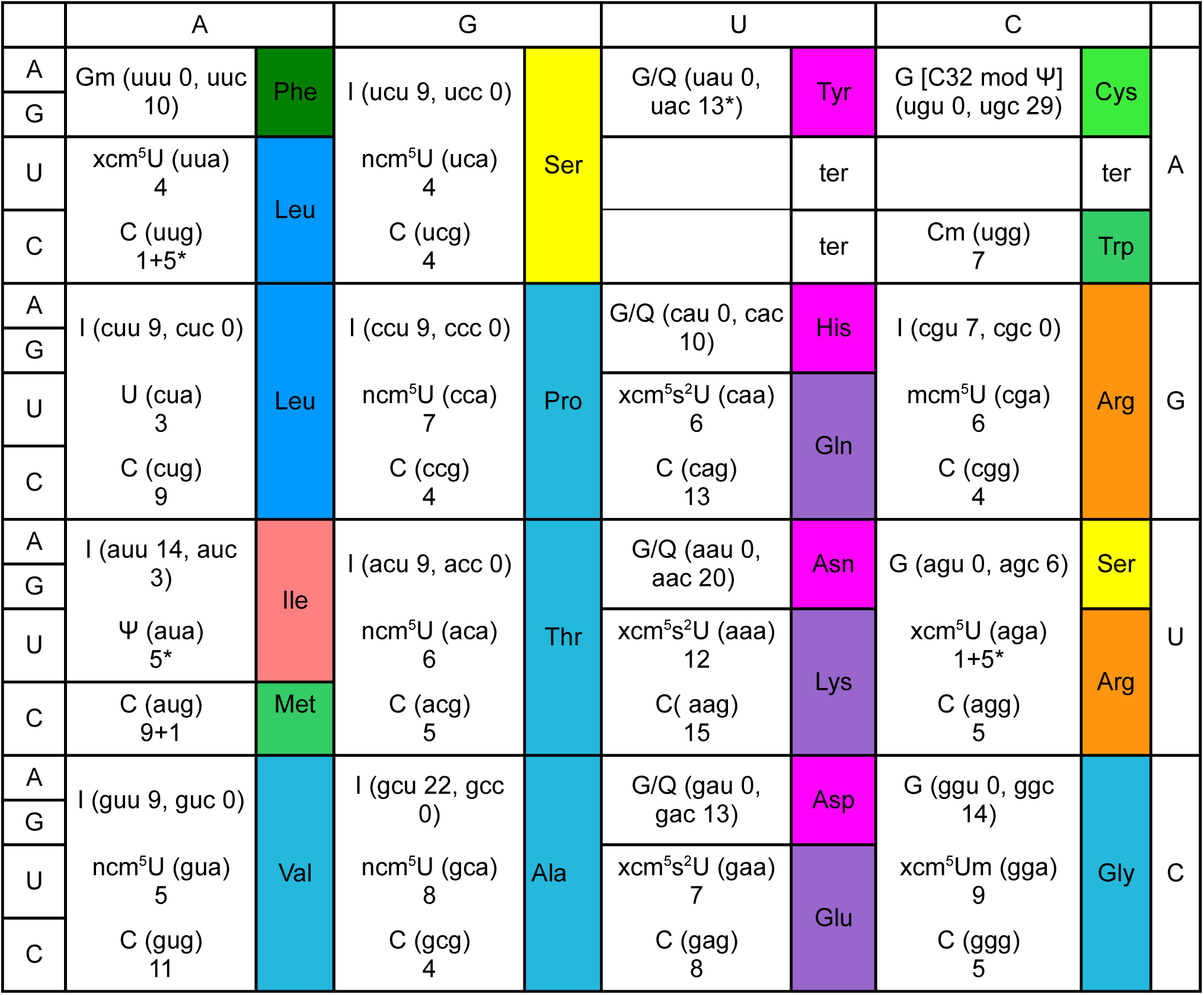
Table of the genetic code with emphasis on decoding by individual tRNAs

The table displays the distribution of the 415 tRNAs coded in the human genome, among which 28 (noted with an asterisk) must splice out an intron as a maturation step: 5 tRNA_Arg_ (AGA), 5 tRNA_Leu_ (UUG), 5 tRNA_Ile_ (AUA) and 13 tRNA_Tyr_ (UAC).

Interestingly, this constraint is easily matched by the tRNA supply of the cell because, contrary to codons ending with a purine, which require distinct tRNAs to be decoded, codons ending with a pyrimidine (U or C, Y) are mainly decoded by a common tRNA species. In the case of NAY codons, the position 34 of tRNAs is a G, replaced by queuine (Q), if this metabolite of bacterial origin is present in the host. Availability of Q in the environment may not have major consequences for translation of NAC codons, but it may alter the speed and accuracy of translation of the NAU codons. G-ending codons are generally rare in the virus, but they do not systematically correspond to rare tRNAs (**Table 2**). Because the anticodon position 34 of the cognate tRNAs is either a guanine or a queuine (Q) residue (depending on a specific input from the environment) and because NAU codons are translated in the absence of Q more slowly and less accurately than NAC codons, a pressure towards NAC in a context where C availability seems to be limiting is probably significant. This may apply to protein Nsp1 and to a lesser extent to protein M (see **Discussion**). All transfer RNA (tRNA) decoding strategies depend on the type and extent of modifications at position 34 of the tRNA anticodon (84). The codon usage bias in SARS-CoV-2 differs from that of average human proteins. In particular it is enriched in codons that require tRNAs modified at position N34 of the anticodon with complex modifications that are linked to zinc homeostasis (85), an important feature knowing that several of the virus functions are Zn^2+^-dependent, while antiviral protein ZAP is a zinc-finger protein (86).

A prominent feature is that two A-ending codons, CUA and CGA are generally rare in the virus sequence (**Figure 5** and **Supplementary Figure 3>**). This is significant, as witnessed by the facts that the cognate codons CUU and CGU are particularly frequent (remember that A and U are both abundant in the virus genome). CUA is decoded by a specific tRNA_Leu_ which is subject to specific regulation (87). In general NNA codons are decoded by tRNAs which have heavily modified U34 residues (88). This imposes a burden on the translation machinery that may impact the viral development, leading to negative selection. An exception is, as reported above, the AGA codon, which is particularly frequent, and this makes protein Nsp1 stand out, highlighting further A-ending codon deficiencies (GGA, CUA, CCA, AGA and GUA). The deficiency of A-ending codons for nucleocapsid protein N, which is very rich in arginine residues and has the expected excess of AGA is limited to UUA, AUA, CUA and GUA, which corresponds to a UpA deficiency in mammalian genomes (89).

While the virus must certainly manage tRNA availability and adapt this resource to its specific codon usage bias, it must also curb the innate antiviral immunity which affects the pool of tRNAs directly. In human cells, tRNA molecules are synthesised as precursors that are maturated into pre-tRNAs that lack their CCA terminal end and need to be further modified (90). Remarkably, stress-induced synthesis of the specific protease angiogenin removes these CCA termini, stopping translation (91). Cells can overcome this process using tRNA nucleotidyl transferase and CTP and ATP. This is yet another CTP-controlled function that must be overcome by the virus. The specificity of angiogenin is modulated by tRNA modifications—this protease can also cut the tRNA molecules at sites located in their anticodons (92)—and this may create an uneven selection pressure on the various tRNAs used to decode the virus genes.

## DISCUSSION

Coronaviruses must tap into their host resources to build up multiple copies of their RNA genome and of their envelope. In the present study we have documented the unique role of CTP as a general coordinator of the cell’s metabolism. Metabolic availability of this nucleotide drives synthesis of the viral genome, its envelope and maintains the translation machinery. This coordinated role makes us understand the presence of the general innate immunity antiviral metabolite, 3’-deoxy-3’,4’-didehydro-CTP (ddhCTP), produced from CTP by the interferon-induced protein viperin (93). Indeed the role of ddhCTP has been somewhat elusive, with experiments suggesting interference with RNA replication/transcription (94), while others demonstrated interference with lipid metabolism (95), and none, at this time, showing that it should impact tRNA synthesis via inhibition of CCAse, a plausible action of this analog of CTP.How did this integrative role of CTP emerge during evolution?

Cells do not have to grow during viral infection. Yet, they result from billion years of evolution based on growth. In a three-dimensional space—the physical space where material entities such as cells flourish—this is akin to squaring the circle. Growth of the cytoplasm (three dimensions) must be matched with growth of the membrane (two dimensions) and growth of the genome (one dimension). Putting together these three facets while sharing a common metabolism cannot be straightforward. Alas, as often in biology, solving a clear functional problem results more often than not in *ad hoc* solutions built on a fairly haphazard collection of bits and pieces, with considerable differences between different species. This would then preclude any consistent view of the anecdotes invented during evolution and end up in a catalog of solutions, as witnessed in the millions of articles that tackle biological questions. We could anticipate that the extensive evolutionary time scale allowed for a slow progression, exploring an infinite variety of directions.

Yet, we could be lucky and discover a universal set-up that may have some generality or even span the whole tree of life. Because the number of the cell’s building blocks is small (mainly nucleotides, amino-acids, phospholipids and carbohydrates), natural selection may have recruited a limited number of those components to implement homeostatic regulation of the growth of the various cell compartments. From detailed analysis of the genome signatures of various organisms and their metabolic constraints, we have demonstrated that the biosynthetic and salvage pathways leading to CTP had remarkable consequences in organizing core cellular functions. A first account of this discovery is presented in (5) and a detailed view is presented in **Figure 2**. The highly involved set-up of this metabolic facet is significant for the manner and process by which a virus invades a cell and subsequently evolves a progressively better adapted progeny. Here we explored, using a functional analysis approach, how understanding this exceptional setup of intermediary metabolism allowed us to anticipate this particular aspect of the evolution of RNA viruses. The main conclusion we reached is that, overall, the coronavirus genomes had a tendency to shed their cytosine —respectively guanine—complement, essentially replacing it by uracil—and guanine by adenine.

While this tendency constrains the genome as a whole, it is obvious that this will dramatically restrict the evolutionary trajectory of the virus, presumably leading to attenuation. However, because a major component of the antiviral innate immunity results from the production of the CTP analog ddhCTP, losing C residues in the genome will transiently alleviate some of the negative pressure created by this antiviral response. This may somehow account for the increase in virulence when a fairly GC-rich virus of an animal infects a foreign host. Furthermore, local C-enrichment, resulting from inevitable template misreading, may be stabilized if the function of the corresponding translated polypeptide contributes to the production of a larger progeny of the virus. This makes identification of the functions associated to loci that do not readily comply with the loss of C (and G) residues as likely candidates important for a stable virus evolutionary potential.

Immediately upon internalization of the virus, its 3’-capped RNA genome begins to be translated into two large proteins coded from ORF1a and ORF1ab which contain a protease domain that cuts off 16 accessory proteins required for specific functions of the virus (96). Its N-terminal domain, processed into non structural protein Nsp1, immediately interferes with translation of the host proteins—it is also inhibiting its own translation thus producing homeostatic regulation—by blocking the assembly of ribosomes that are in the process of translating host mRNAs and disrupting nuclear-cytoplasmic transport (97). Subsequently a large complex forms with all the other Nsp proteins generated from the processed precursor, generating a RNA-dependent replication/transcription complex (RTC). Remarkably, this complex is tightly linked to key elements of the translation machinery. It has been demonstrated that, besides inhibition of interferon signalling, Nsp1 binds to the 40S ribosomal subunit (98) and that it further triggers host mRNA degradation (99). Nsp1 binds translation factors eIF3, eIF1A, eIF1 and eIF2-tRNAi-GTP (100) and inhibits formation of the translation initiation complex—48S complex and formation of active 80S ribosomes (101). Early interaction with the host translation machinery will stop translation, triggering host mRNA decay, which both produces nucleotide precursors for replication of the virus and hijacks the machinery to perform further translation of the viral genome. This implies that the complex between Nsp1 and the translation initiation complex is able to discriminate between different classes of mRNAs to allow or prevent their translation. Among the factors bound to Nsp1 the enigmatic ATP-dependent enzyme ABCE1 has been identified (100). Remarkably, this protein is expected to behave as a “Maxwell’s demon” as do proteins of the EttA family in Bacteria, allowing partition of specific mRNA families the expression of which needs to be co-expressed or co-repressed (102).

This role of translation is apparent in the codon usage bias of Nsp1, which differs from that of subsequent domains cleaved off ORF1a and ORF1ab polypeptides. Here the role of arginine residue codons seems to have been submitted to strong selection, with a majority being CGU codons, while AGA and AGG codons— AGA translation being over-represented in the subsequent polypeptides—are totally absent from the sequence. Also, the arginine codon CGG is extremely rare overall in the genome sequence, and its locations are revealing, likely to be important for the co-translational folding of cognate proteins. This is particularly important in SARS-CoV-2 as the insertion generating its furin-like cleavage site in the spike protein that mediates cell entry (103–105) is located right at a CGG doublet. Besides protein Nsp1, we demonstrated that the nucleocapsid protein N had also a general distribution of the codon usage bias that differed from that of the bulk of proteins coded from the ORF1a and ORF1ab regions. This is likely to be due to the fact that the corresponding transcripts also code for another protein in a different reading frame, protein ORF9b (106), and we can assume that this observations substantiates that this protein has indeed an important role in the biology of the virus. Remarkably, this also makes that the codon and tRNA usage bias of protein N resembles that of the human host. Whether this is meaningful should be further explored.

## PERSPECTIVES

Here, we reviewed the role of a specific intracellular metabolic pressure that must constrain the evolution of the genome sequence of RNA viruses, with emphasis on SARS-CoV-2 and in the context of the entire coronavirus family. Several studies have noticed the cytosine deficiency in the genome of evolving coronaviruses, with concomitant deficiency in the position of codons, but these observations were ascribed to deamination of cytosine resulting from the action of the host APOBEC system (107) or to methylation of CpG dinucleotides (108) as driving forces for evolution. By contrast, our working hypothesis is that the availability of CTP (and hence of cytosine-based precursors) is a dominating driving force in the way the virus evolves a new progeny. We are well aware that, due to the small number of samples and fairly short life time (as compared to usual evolutionary trajectory of a virus species), sequence analyses would provide only a limited view of sequence evolution and should be used more as a “rule of thumb” than a view based on trustworthy statistics. However we believe that, in view of the urgent situation we are facing, it is important to communicate our observations while relating them to previously unrecognised pressure that must have considerable importance in the evolution of viruses and the metabolic backdrop of its biology in the host cell. In this respect it seems worthwhile to explore whether unappreciated functions coded by viruses will be involved in controlling CTP availability.

## MATERIAL AND METHODS

### Data preparation

Dataset of reference coronaviruses: viral genomes were downloaded from GenBank (https://www.ncbi.nlm.nih.gov/genbank/) and GISAID (https://www.gisaid.org/). Representative coronaviruses of different species were selected from complete genomes, with reference genomes recommended by the Coronaviridae Study Group of the International Committee on Taxonomy of Viruses (https://talk.ictvonline.org/) and NCBI retained preferentially. For viruses containing isolates from different hosts, at least one representative strain from each host was kept. The genomes were aligned using MAFFT v7.427 (109) and manually checked with BioEdit. Alignment of full genomic sequences were used for phylogeny reconstruction, while the coding regions for ORF1ab were extracted for codon usage analysis.

Dataset of SARS-CoV-2: a total of 17037 SARS-CoV-2 related sequences were available from GISAID on May 6^th^, 2020 (110). Only SARS-CoV-2 genomes isolated from human, with a full length over 27,000 bp, no ambiguous sites and detailed collection date information were used for alignment. For duplicate sequences, only the earliest isolate was kept. Sequences for 26 coding regions, including Nsp1, Nsp2, Nsp3, Nsp4, Nsp5, Nsp6, Nsp7, Nsp8, Nsp9, Nsp10, Nsp11, Nsp12, Nsp13, Nsp14, Nsp15, Nsp16, S, ORF3a, E, M, ORF6, ORF7a, ORF7b, ORF8, N and ORF10 were extracted for each strain, using NC_045512 as reference. The coding sequences were checked manually to exclude those with abnormal mutations and early stop codons. A total of 4,110 strains with all 26 coding regions of complete ORF length were retained. After further deduplication based on the concatenated sequences comprised of the 26 ORFs, the final dataset contained a total of 2,574 unique SARS-CoV-2 isolates.

### Phylogeny reconstruction

Phylogenetic tree of the 89 representative coronaviruses was inferred using the Maximum Likelihood (ML) method implemented in IQ-TREE v1.6.12 with the GTR+F+I+G4 substitution model determined by ModelFinder (111–113). Ultra-fast bootstrap support values were calculated from 1,000 pseudo-replicate trees (112). Visualization of phylogenies were conducted with ggtree package (114).

### Gap-based alignment

The full alignment of the 89 reference strains was used to generate a tree, using FastTree 2.1.10 (115) (with gamma distribution and the nucleotide option on – namely with the command options -gamma -nt), on the NGPhylogeny.fr server (116). The Jukes-Cantor model with balanced support Shimodaira-Hasegawa test was selected (117). Total branch length was: 14.267.

Furthermore, a gap-based alignment was created, using gaps as follows: all dinucleotides were replaced with the ‘undefined’ symbol ‘x’ and the ‘dummy’ symbols (W for 3, Y for 6 and F for 9 consecutive gaps and the V symbol for all single gaps), leaving only single-nucleotides in-between gaps as anchor points (7% of total). The encoding in gaps of 3/6/9 are use to emulate the importance of potential codon gaps (reflected in the BLOSUM45 matrix). Total branch length was: 1.673.

Gap-based genome-based phylogenetic reconstruction for this group is based on the fact that, as also mentioned recently elsewhere (118), these viruses undergo significant recombination and a large number of nucleotide positions achieve saturation thus confounding phylogenetic signal. Tree visualization was facilitated by IcyTree (119)

### Base content calculation

Base content was calculated by dividing the occurrence of each base by the total length of the sequence. Genomic base contents of representative coronaviruses were calculated with the full viral genome sequences. For the base content dynamic analysis of SARS-CoV-2, base compositions were calculated using the 2754 unique sequences concatenated by 26 ORFs.

### Codon usage analysis

Codon usage analysis was conducted based on the ORF1ab region of representative coronaviruses and the 26 individual ORFs of SARS-CoV-2 strains. Relative synonymous codon usage (RSCU) value was defined as the ratio of the observed codon usage to the expected value (120). Codons with an RSCU value of 0, 0∼0.6, 0.6∼1.6 or > 1.6 were regarded as not-used, under-represented, normally-used, or over-represented (121). RSCUs for the 120,426 human coding regions were determined based on the *Homo sapiens* codon usage table retrieved on 2020 June 14^th^ from TissueCoCoPUTs (122).

### Statistical analysis and plots

Statistical test, linear regression and data visualization were all conducted in R. Kruskal-Wallis test by rank and Wilcoxon rank sum test for pairwise comparisons were applied as appropriate. P values are labeled as follows: < 0.0001, ****; 0.0001 to 0.001, ***; 0.001 to 0.01, **; 0.01 to 0.05, *; ≥ 0.05, not labeled. p<0.05 was considered as significant.

## Supporting information

Supplementary Table 2 GISAID Acknowledged

## ACKNOWLEDGEMENTS

We warmly thank Prof. Huanming Yang, Dr. Ziqing Deng and Dr. Minfeng Xiao for their constructive communication in study design and data interpretation. Our thanks also go to Mr. Jielun Cai and Miss Xinyi Cheng for their assistance in data preparation. Thanks also to Pierre-Yves Bourguignon for his analysis of the cytosine complement of the virus and to Agnieszka Sekowska for her comments on metabolic pathways. We thank all the authors who shared genomic data in public database, and an acknowledgment table for sequences retrieved from GISAID (110) is provided (**Supplementary Table S2**).

## CONTRIBUTIONS

AD designed the study and wrote the bulk of the manuscript. ZO, JL and WC developed the in silico analyses of cytosine evolution, codon usage biases and phylogeny. ZO, DW and WS performed the analysis and ZO wrote the corresponding part of the manuscript. CO designed and performed phylogenetic studies based on indels. PM identified the importance of several key steps in CTP synthesis. All authors read, wrote some sections and contributed to the final version of the manuscript.

## SUPPLEMENTARY MATERIAL

**Supplementary Table S1. List of representative coronaviruses for phylogeny reconstruction and codon usage analysis with ORF1ab**

**Supplementary Table S2. Acknowledgement Table for the 2**,**574 SARS-CoV-2 isolates retrieved from GISAID (www.gisaid.org)**

**Supplementary Figure 1: Phylogeny and nucleotide composition of representative coronaviruses**. The Maximum Likelihood phylogeny (left panel) was reconstructed based on the complete genomic sequences. Bootstrap values over 70 are displayed on nodes. Coronaviruses that infects human are highlighted by a red dot at tip. Genus of each strain is indicated by the prefix of the taxa name. The nucleotide composition (right panel) was calculated based on the full genome sequences.

**Supplementary Figure 2: Correlation between base content of SARS-CoV-2 coding regions**. The calculation was based on 2,574 unique SARS-CoV-2 coding sequences concatenated by 26 ORFs.

**Supplementary Figure 3: Details of the codon usage bias of SARS-CoV-2 ORFS viewed from the human tRNA complement**. The X-axis displays codons sorted by amino acid and then by tRNA usage. The Y-axis indicates the mean RSCU value determined for each ORF or the human total coding sequences. Codons using the same tRNA are labeled with the same color.

**Supplementary Figure 4: Phylogeny and codon usage of representative coronaviruses**

A Maximum Likelihood phylogeny (left panel) was reconstructed based on complete genomic sequences of coronaviruses. Bootstrap values over 70 are displayed on nodes. Coronaviruses that infect human are highlighted by a red dot. The genus of each strain is indicated by the prefix of the taxon name. The codon usage (right panel) was calculated based on the ORF1ab region as recorded at GenBank, except for those with incomplete annotations. The X-axis displays codons sorted by the third position, with the first letter denoting the amino acid and the three letters suffix following a dot representing the codon. Codons that are under-represented (RSCU<0.6), normally utilized (RSCU ranges between 0.6-1.6), or over-represented (RSCU>1.6) are labeled in blue, ice cold and yellow, respectively.

