## Supplementary Tables and Figures for "A path towards SARS-CoV-2 attenuation: metabolic pressure on CTP synthesis rules the virus evolution"

### Supplementary Table S1. List of representative coronaviruses for phylogeny reconstruction and codon usage analysis with ORF1ab

| Genus | Virus | Host | Length (bp) | GenBank |
|---|---|---|---|---|
| AlphaCoV | Alpaca respiratory coronavirus CA08-1/2008 | Alpaca | 27374 | JQ410000 |
| AlphaCoV | Bat coronavirus BtKYNL63-9a | Bat | 28363 | NC_032107 |
| AlphaCoV | Bat coronavirus BtKY229E-1 | Bat | 27837 | KY073747 |
| AlphaCoV | Bat coronavirus BtKYNL63-9b | Bat | 28679 | KY073745 |
| AlphaCoV | Bat coronavirus HuB2013 | Bat | 28755 | KJ473798 |
| AlphaCoV | Bat coronavirus 1A | Bat | 28326 | NC_010437 |
| AlphaCoV | Bat coronavirus 008_16/M.bra/FIN/2016 | Bat | 28119 | MN065811 |
| AlphaCoV | Bat coronavirus CDPHE15 | Bat | 28035 | NC_022103 |
| AlphaCoV | BatMr coronavirus SAX2011 | Bat | 27935 | NC_028811 |
| AlphaCoV | BatNv coronavirus SC2013 | Bat | 27783 | NC_028833 |
| AlphaCoV | BatRf coronavirus YN2012 | Bat | 26975 | NC_028824 |
| AlphaCoV | BatRf coronavirus HuB2013 | Bat | 27608 | NC_028814 |
| AlphaCoV | Camel coronavirus Riyadh/Ry141/2015 | Camel | 27395 | NC_028752 |
| AlphaCoV | Canine coronavirus 1 strain 23/03 | Canine | 30004 | KP849472 |
| AlphaCoV | Canine coronavirus strain 1-71 | Canine | 29462 | JQ404409 |
| AlphaCoV | Feline alphacoronavirus 1 SB22 | Feline | 29137 | MH817484 |
| AlphaCoV | Feline infectious peritonitis virus 79-1146 | Feline | 29355 | NC_002306 |
| AlphaCoV | Ferret coronavirus FRCoV-NL-2010 | Ferret | 28434 | NC_030292 |
| AlphaCoV | Ferret coronavirus FRCoV4370 | Ferret | 28550 | LC119077 |
| AlphaCoV | Hipposideros pomona bat coronavirus CHB25 | Hipposideros | 28169 | MN611525 |
| AlphaCoV | Human coronavirus NL63 | Human | 27553 | NC_005831 |
| AlphaCoV | Human coronavirus 229E | Human | 27317 | NC_002645 |
| AlphaCoV | Miniopterus bat coronavirus HKU8 | Miniopterus | 28773 | NC_010438 |
| AlphaCoV | Mink coronavirus WD1127 | Mink | 28941 | NC_023760 |
| AlphaCoV | Mink coronavirus China/1/2016 | Mink | 28924 | MF113046 |
| AlphaCoV | Murine coronavirus AcCoV-JC34 | Murine | 27682 | NC_034972 |
| AlphaCoV | Porcine epidemic diarrhea virus | Porcine | 28033 | NC_003436 |
| AlphaCoV | Porcine transmissible gastroenteritis virus AHHF | Porcine | 28614 | KX499468 |
| AlphaCoV | Rat coronavirus Lucheng-19 | Rat | 28763 | KF294380 |
| AlphaCoV | Rhinolophus bat coronavirus HKU32 | Rhinolophus | 29201 | MK720946 |
| AlphaCoV | Rhinolophus bat coronavirus HKU2 | Rhinolophus | 27165 | NC_009988 |
| AlphaCoV | Rousettus bat coronavirus HKU10-183A | Rousettus | 28494 | NC_018871 |
| AlphaCoV | Scotophilus bat coronavirus 512 | Scotophilus | 28203 | NC_009657 |
| AlphaCoV | Swine acute diarrhea syndrome | Swine | 27184 | MH615810 |
| AlphaCoV | Swine enteric coronavirus | Swine | 28111 | NC_028806 |
| AlphaCoV | Tylonycteris bat coronavirus HKU33 strain [illegible] | Tylonycteris | 27636 | MK720944 |
| BetaCoV | Bat coronavirus BM48-31/BGR/2008 | Bat | 29276 | NC_014470 |
| BetaCoV | Bat coronavirus HKU9-2 | Bat | 29107 | EF065514 |
| BetaCoV | Bat coronavirus SC2013 | Bat | 30423 | KJ473821 |
| BetaCoV | Bat coronavirus RaTG13 | Bat | 29855 | MN996532 |
| BetaCoV | Bat coronavirus HKU3-1 | Bat | 29728 | DQ022305 |
| BetaCoV | Bat coronavirus WIV16 | Bat | 30290 | KT444582 |
| BetaCoV | Bat coronavirus JPDB144 | Bat | 30321 | KU182965 |
| BetaCoV | Bat coronavirus PREDICT/PDF-2180 | Bat | 29642 | NC_034440 |

| Genus | Virus | Host | Length (bp) | GenBank |
|---|---|---|---|---|
| BetaCoV | Bat coronavirus Rp3/2004 | Bat | 29736 | DQ071615 |
| BetaCoV | Bat coronavirus Yunnan2011 | Bat | 29452 | JX993988 |
| BetaCoV | BatHp coronavirus Zhejiang2013 | Bat | 31491 | NC_025217 |
| BetaCoV | Bovine coronavirus ENT | Bovine | 31028 | NC_003045 |
| BetaCoV | Canine respiratory coronavirus K37 | Canine | 31028 | JX860640 |
| BetaCoV | Civet coronavirus SZ3/2003 | Civet | 29741 | AY304486 |
| BetaCoV | Dromedary camel coronavirus HKU23 | Dromedary | 31075 | MN514966 |
| BetaCoV | Equine coronavirus NC99 | Equine | 30992 | EF446615 |
| BetaCoV | Giraffe coronavirus US/OH3/2003 | Giraffe | 31002 | EF424623 |
| BetaCoV | Hedgehog coronavirus | Hedgehog | 30148 | NC_039207 |
| BetaCoV | Human coronavirus HKU1 | Human | 29926 | NC_006577 |
| BetaCoV | Human coronavirus OC43 strain | Human | 30741 | NC_006213 |
| BetaCoV | Human SARS coronavirus Tor2 | Human | 29751 | NC_004718 |
| BetaCoV | Human SARS-CoV-2 strain Wuhan- | Human | 29903 | NC_045512 |
| BetaCoV | Human MERS coronavirus | Human | 30119 | NC_019843 |
| BetaCoV | Hypsugo bat coronavirus HKU25 isolate YD131305 | Hypsugo bat | 30498 | KX442564 |
| BetaCoV | Murine coronavirus MHV-3 | Murine | 31448 | FJ647224 |
| BetaCoV | Pangolin coronavirus PCoV_GX-P5L | Pangolin | 29806 | MT040335 |
| BetaCoV | Pipistrellus bat coronavirus HKU5 | Pipistrellus | 30482 | NC_009020 |
| BetaCoV | Porcine hemagglutinating encephalomyelitis JL/2008 | Porcine | 30684 | KY994645 |
| BetaCoV | Rabbit coronavirus HKU14 | Rabbit | 31100 | NC_017083 |
| BetaCoV | Rat coronavirus Parker | Rat | 31250 | NC_012936 |
| BetaCoV | Rat Betacoronavirus HKU24 | Rat | 31249 | NC_026011 |
| BetaCoV | Rousettus bat coronavirus HKU9 | Rousettus | 29114 | NC_009021 |
| BetaCoV | Rousettus bat coronavirus GCCDC1 356 | Rousettus | 30161 | NC_030886 |
| BetaCoV | Sable antelope coronavirus US/OH1/2003 | Sable | 30995 | EF424621 |
| BetaCoV | Tylonycteris bat coronavirus HKU4 | Tylonycteris | 30286 | NC_009019 |
| BetaCoV | Water deer coronavirus W17-18 | Water deer | 31034 | MG518518 |
| BetaCoV | Yak coronavirus YAK/HY24/CH/2017 | Yak | 31032 | MH810163 |
| DeltaCoV | Bulbul coronavirus HKU11-934 | Bulbul | 26487 | NC_011547 |
| DeltaCoV | Common moorhen coronavirus HKU21 | Common | 26223 | NC_016996 |
| DeltaCoV | Magpie-robin coronavirus HKU18 | Magpie-robin | 26689 | NC_016993 |
| DeltaCoV | Munia coronavirus HKU13-3514 | Munia | 26552 | NC_011550 |
| DeltaCoV | Night heron coronavirus HKU19 | Night | 26077 | NC_016994 |
| DeltaCoV | Porcine coronavirus HKU15-155 | Porcine | 25425 | NC_039208 |
| DeltaCoV | Sparrow coronavirus HKU17 | Sparrow | 26083 | NC_016992 |
| DeltaCoV | Thrush coronavirus HKU12-600 | Thrush | 26396 | NC_011549 |
| DeltaCoV | White-eye coronavirus HKU16 | White-eye | 26041 | NC_016991 |
| DeltaCoV | Wigeon coronavirus HKU20 | Wigeon | 26227 | NC_016995 |
| GammaCoV | Avian infectious bronchitis virus | Avian | 27608 | NC_001451 |
| GammaCoV | Beluga whale coronavirus SW1 | Beluga | 31686 | NC_010646 |
| GammaCoV | Bottlenose dolphin coronavirus HKU22 strain | Bottlenose | 31769 | KF793824 |
| GammaCoV | Duck coronavirus DK/GD/27/2014 | Duck | 27754 | KM454473 |
| GammaCoV | Pheasant coronavirus ph/China/I0710/17 | Pheasant | 27655 | MK423876 |
| GammaCoV | Turkey coronavirus MG10 | Turkey | 27657 | NC_010800 |

**Supplementary Table S2. Acknowledgement Table for the 2,574 SARS-CoV-2 isolates retrieved from GISAID (www.gisaid.org)**

Excel Table

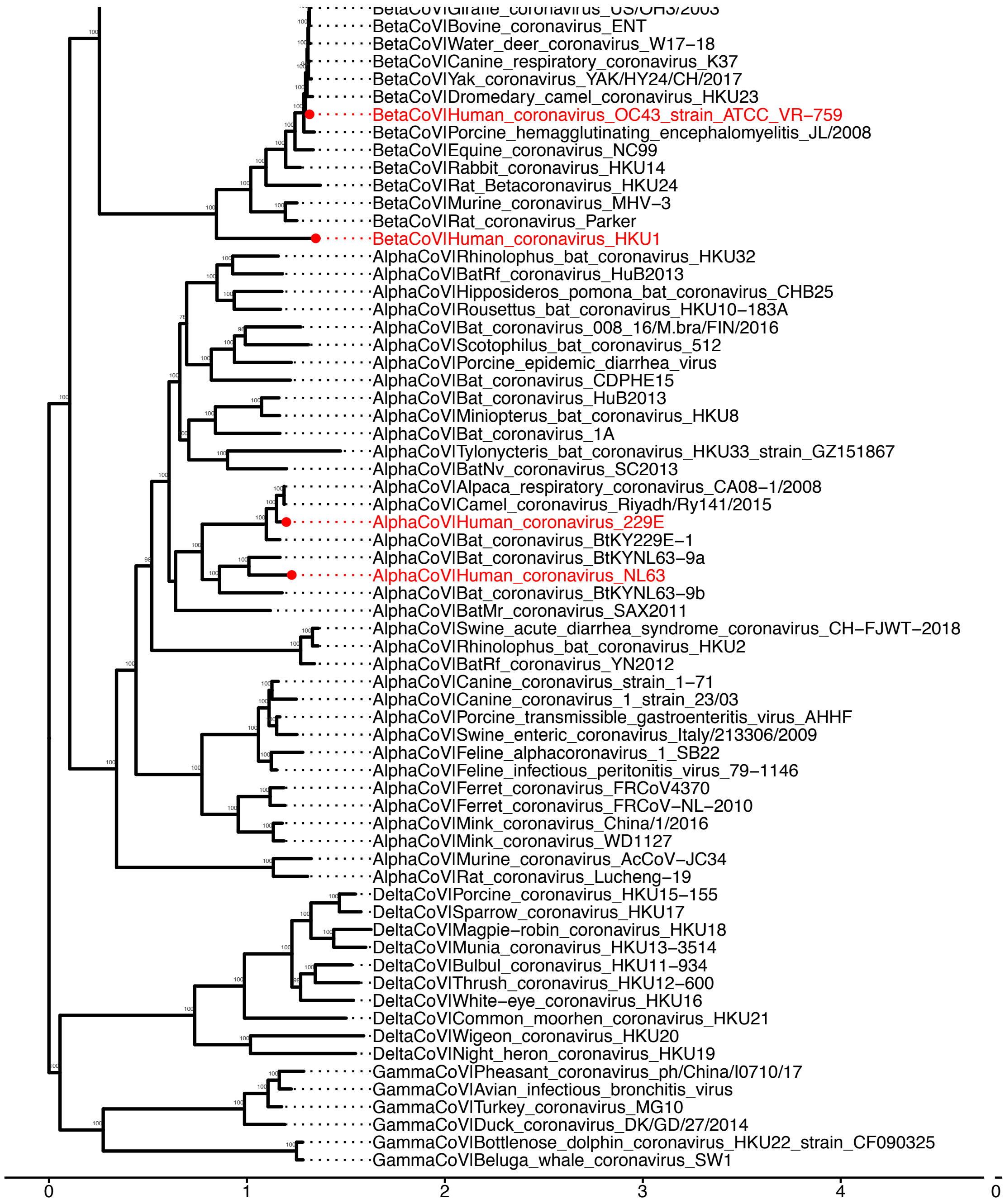


**Supplementary Figure 1: Phylogeny and nucleotide composition of representative coronaviruses.** The Maximum Likelihood phylogeny (left panel) was reconstructed based on the complete genomic sequences. Bootstrap values over 70 are displayed on nodes. Coronaviruses that infects human are highlighted by a red dot at tip. Genus of each strain is indicated by the prefix of the taxa name. The nucleotide composition (right panel) was calculated based on the full genome sequences.

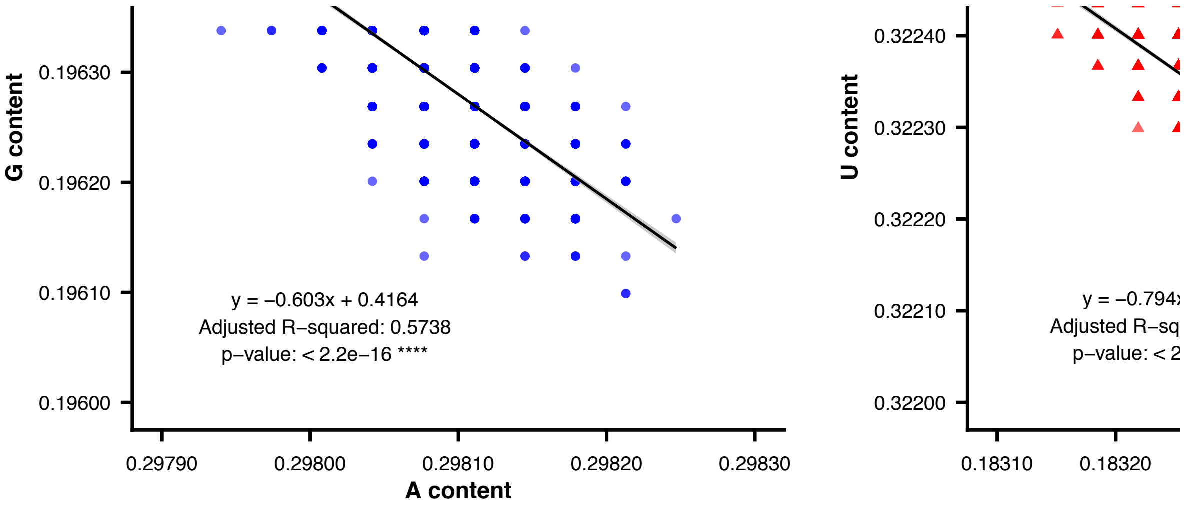


**Supplementary Figure 2: Correlation between base content of SARS-CoV-2 coding regions.** The calculation was based on 2574 unique SARS-CoV-2 coding sequences concatenated by 26 ORFs.

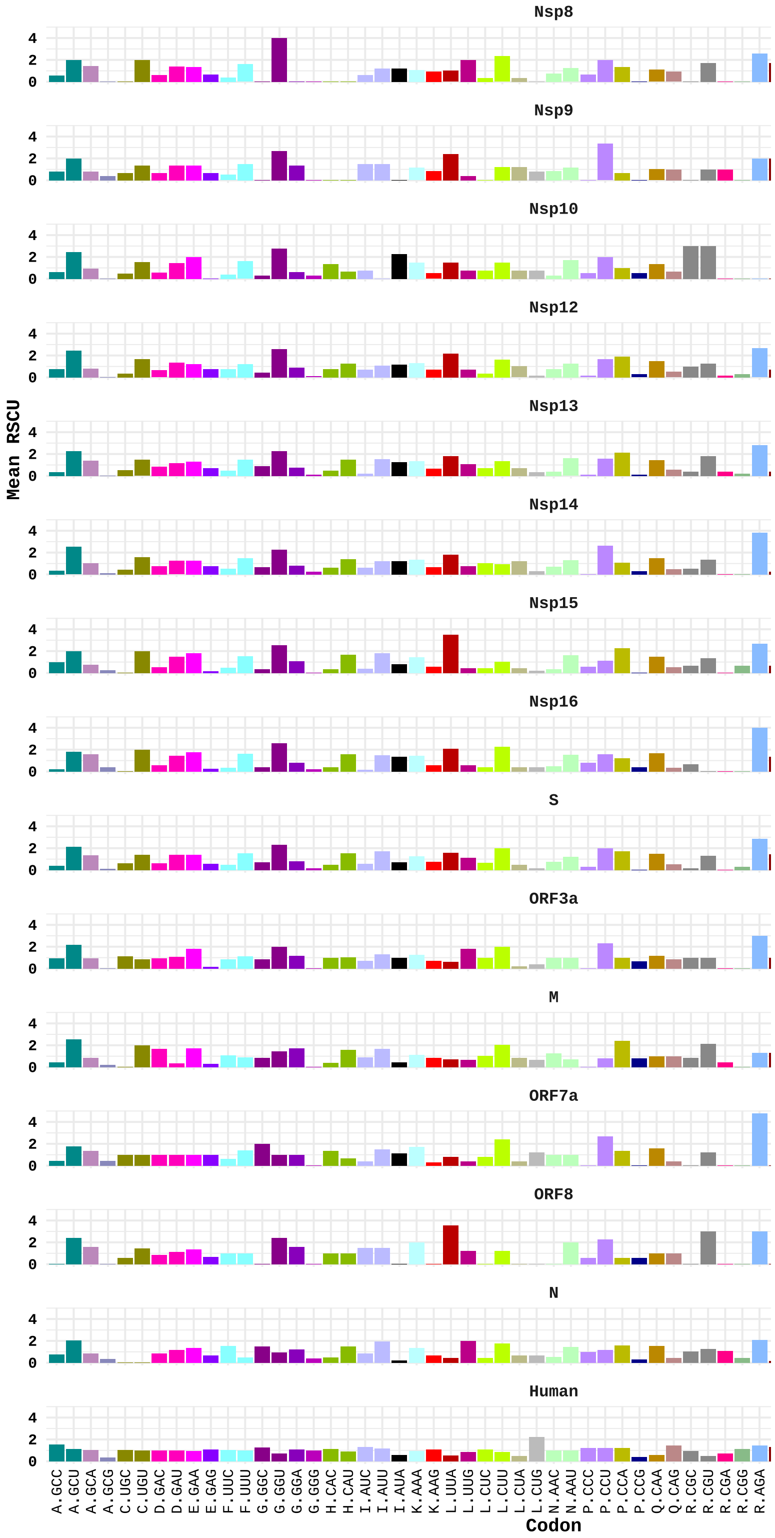


**Supplementary Figure 3: Details of the codon usage bias of SARS-CoV-2 ORFS viewed from the human tRNA complement.** The X-axis displays codons sorted by amino acid and then by tRNA usage. The Y-axis indicates the mean RSCU value determined for each ORF or the human total coding sequences. Codons using the same tRNA are labeled with the same color.

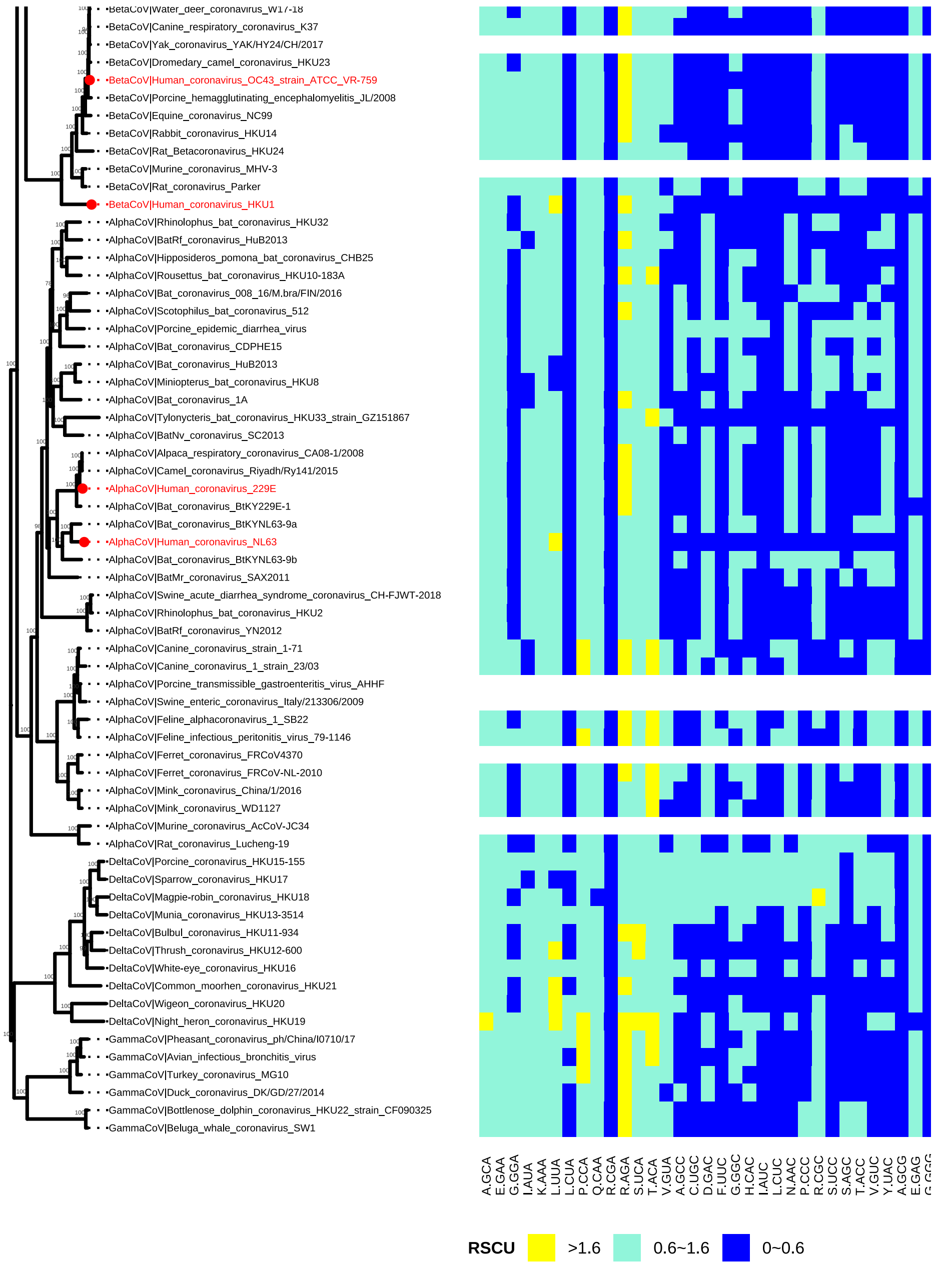


**Supplementary Figure 4: Phylogeny and codon usage of representative coronaviruses.** The Maximum Likelihood phylogeny (left panel) was reconstructed based on the complete genomic sequences. Bootstrap values over 70 are displayed on nodes. Coronaviruses that infects human are highlighted by a red dot at tip. Genus of each strain is indicated by the prefix of the taxa name. The codon usage (right panel) was calculated based on the ORF1ab region as recorded in NCBI, except for those with incomplete annotations. The X-axis displays codons sorted by the third position, with the first letter denoting the amino acid and the three letters suffix the dot representing the codon. Codons that are under-represented (RSCU<0.6), normally utilized (RSCU ranges between 0.6-1.6), or over-represented (RSCU>1.6) are labeled in blue, ice cold and yellow, respectively.
